# Mapping multicellular programs from single-cell profiles

**DOI:** 10.1101/2020.08.11.245472

**Authors:** Livnat Jerby-Arnon, Aviv Regev

**Affiliations:** Klarman Cell Observatory, Broad Institute of MIT and Harvard, Cambridge, MA; Howard Hughes Medical Institute and Koch Institute of Integrative Cancer Research, Department of Biology, Massachusetts Institute of Technology, Cambridge, MA; Genentech, 1 DNA Way, South San Francisco, CA

## Abstract

Tissue homeostasis relies on orchestrated multicellular circuits, where interactions between different cell types dynamically balance tissue function. While single-cell genomics identifies tissues’ cellular components, deciphering their coordinated action remains a major challenge. Here, we tackle this problem through a new framework of multicellular programs: combinations of distinct cellular programs in different cell types that are coordinated *together* in the tissue, thus forming a higher order functional unit at the tissue, rather than only cell, level. We develop the open-access DIALOGUE algorithm to systematically uncover such multi-cellular programs not only from spatial data, but even from tissue dissociated and profiled as single cells, *e.g.*, by single-cell RNA-Seq. Tested on spatial transcriptomes from the mouse hypothalamus, DIALOGUE recovered spatial information, predicted the properties of a cell’s environment only based on its transcriptome, and identified multicellular programs that mark animal behavior. Applied to brain samples and colon biopsies profiled by scRNA-Seq, DIALOGUE identified multicellular configurations that mark Alzheimer’s disease and ulcerative colitis (UC), including a program spanning five cell types that is predictive of response to anti-TNF therapy in UC patients and enriched for UC risk genes from GWAS, each acting in different cell types, but all cells acting in concert. Taken together, our study provides a novel conceptual and methodological framework to unravel multicellular regulation in health and disease.

## INTRODUCTION

The interplay between different cells in a tissue ecosystem is crucial for maintaining tissue homeostasis. While many diseases have been traditionally perceived as the result of the malfunction of a particular cell or cell type, mounting evidence (1–5) and new therapeutic strategies (3,6,7) have demonstrated the pivotal role of multicellular action in maintaining homeostasis and its dysregulation in a wide-range of diseases, thus opening new opportunities for intervention, diagnosis, disease monitoring and prevention. In parallel, advances in single cell RNA-seq (scRNA-seq) (8,9) and spatial transcriptomics (10–12) now allow us to systematically explore molecular profiles at single cell resolution across cell types (13), tissues (14,15), and disease states (16–18), in both isolated cells and intact tissues (10–12). However, despite these advances, deciphering multicellular regulation still remains a major challenge, limiting our ability to move from a cell- to a tissue-centric perspective.

One of the key current limitations is in our computational formulation of the problem and available analysis methods. While many computational methods have been developed to analyze single cell data the vast majority were geared to map and explore single cell states (13,19–25). Those developed to study cell-cell interactions, are focused on reconstructing the tissue’s spatial organization (*e.g.*, by combining single cell and spatial data or based on assumptions on spatial patterning) (26–29), on inferring putative physical cell-cell interactions based on known receptor-ligand pairs and known signaling pathways (30–32), or on highlighting recurring cell type compositions of cellular neighborhoods using multiplex molecular spatial data (33,34). Thus, while these methods revealed important properties of cell biology and tissue structure, we still lack a computational basis to study tissue biology and functional multicellular processes at scale.

Here, we approach this problem in a new way, by introducing the concept of multicellular programs and developing the first algorithm to systematically uncover multicellular programs from single-cell data. We define multi-cellular programs as the combinations of different cellular programs in different cell types that are coordinated together in the tissue, thus forming a higher-order functional unit at the tissue-level, rather than only at the cell-level. We develop DIALOGUE, a computational approach for <u>d</u>ecoupling cell st<u>a</u>tes through multicellular c<u>o</u>nfi<u>gu</u>ration id<u>e</u>ntification, by using cross-cell-type associations to identify multicellular programs and map the cell transcriptome as a function of its environment. We apply DIALOGUE to a wide-range single-cell data types, obtained not only from spatially intact tissue (*e.g.*, MERFISH) but even from dissociated cells (*e.g.*, single-cell RNA-Seq). Tested on MERFISH data from the mouse hypothalamus (10) DIALOGUE recovered spatial information, predicted the properties of a cell’s environment only based on its transcriptome, and identified multicellular programs that mark animal behavior. Next, applied to single-cell RNA-seq from colon biopsies, DIALOGUE identified a coordinated multicellular response across epithelial, innate and adaptive immune cells that arises in ulcerative colitis patients, predicts clinical responses to anti-TNF therapy, and spans multiple genes associated with UC risk by Genome Wide Association Studies (GWAS) but acting in different cells. Lastly, using single nucleus transcriptomes from the prefrontal cortex, DIALOGUE highlighted an Alzheimer’s disease multicellular program that increases in the aging brain. Taken together, our framework and the DIALOGUE method provide a systematic way to study cellular crosstalk, along with its molecular underpinnings and phenotypic implications, as part of a transition from single cell to tissue biology.

## RESULTS

### DIALOGUE: <u>D</u>ecoupling cell st<u>a</u>tes through multicellular c<u>o</u>nfi<u>gu</u>ration id<u>e</u>ntification

DIALOGUE leverages the reasoning that cells within the same microenvironment are exposed to similar cues, which may activate coordinated responses in different cell types at two (non-mutually exclusive) levels (**Fig. 1A**). First, cells of multiple types may simultaneously activate the same, cell-type-independent program (35) (**Fig. 1A**). Second, cells of different types may each activate a different, cell-type-specific program, but in a concerted fashion, either because they directly impact each other, or because they each respond distinctly to the same shared cue (**Fig. 1A**). Both cases should give rise to corresponding transcriptional programs across different cell types – where the expression of one set of genes in a certain cell is associated with the expression of the same or another set of genes in nearby cells or cells from the same sample. If we can find those patterns from data, we should be able to uncover multicellular configurations and their dependent cellular programs.

**Figure 1.**
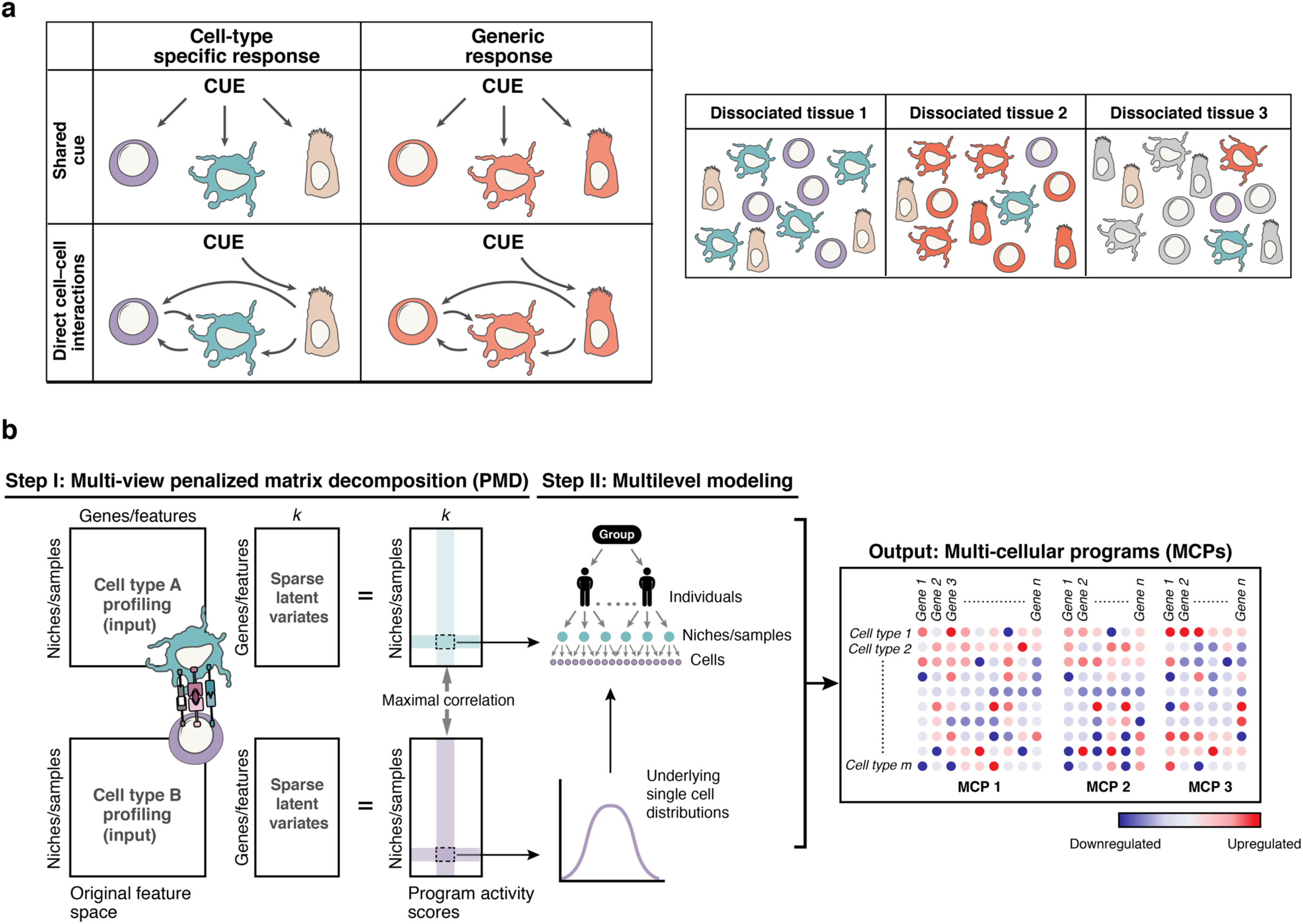
DIALOGUE: a dimensionality reduction method for uncovering multicellular programs from single-cell data. **(A)** Multicellular programs (MCPs). Right: Multicellular programs can arise due to either shared cues (upper row) or direct cell-cell interactions (bottom row) and comprise both cell type specific (left column) or shared (right column) programs. Left: Multicellular programs can be observed also in dissociated tissue sections, as their different cell-type-specific and generic components are statistically dependent across different cell types. (B) DIALOGUE method. In Step I (left), DIALOGUE takes as input (left) mean gene/feature expression (columns) matrices for each cell type, across samples (e.g., scRNA-seq data) or physical niches (*in situ* data) (rows) and infers latent features (“sparse latent variates”) across those samples/niches (middle matrices) and their activity (right matrices) for each cell type, such that the programs of each cell type are highly correlated with the corresponding programs of all the other cell types (in this case shown only for two). In Step II (right), it identifies a gene signature for each sparse canonical variate, by interrogating the single-cell distributions, while accounting for potential confounding factors at different levels. This results in a representation of multicellular programs, each with a set of up- and down-regulated genes in each of its cell type compartments (right).

Given single-cell profiles from different spatial locations (in spatial profiling) or from different samples (in scRNA-seq), DIALOGUE treats different types of cells from the same micro/macroenvironment or sample as different representations of the same entity (**Fig. 1B**). It first applies penalized matrix decomposition (PMD) (36) to identify sparse canonical variates that transform the original features’ space (*e.g.*, genes, PCs, etc.) to a new feature space, where the different (cell-type-specific) representations are correlated across the different samples/environments; this is done based on the average signal across the different cell subsets in each niche/sample. It then identifies the specific genes that comprise these latent features by fitting multilevel (hierarchical) models; this step models the single-cell distributions and controls for potential confounders, such as gender, age, variation in sample preparation, technical variability etc. (**Fig. 1B, Methods**). As output, DIALOGUE provides multicellular programs, each composed of two or more co-regulated, cell-type-associated programs (**Fig. 1B, Methods**).

DIALOGUE can identify numerous multicellular programs (MCPs; input parameter *k* < rank of the original feature space). In practice, the different MCPs are not correlated with one another (**Methods**), and the cross-cell-type correlations observed within an MCP decreases with *k*, such that the first few MCPs depict most of the multicellular co-expression. Importantly, DIALOGUE does not depend on *k* and will find the same MCPs or a subset of them.

Using the multicellular programs, DIALOGUE can accurately identify mis-localized cells and predict the cell’s environment solely based on its transcriptional state (without using any type of spatial information for training). It can decouple cell states to their niche/sample-dependent and independent programs, and further decompose the niche/sample-dependent programs into generic (shared) and cell-type-specific components (**Fig. 1A,B**). DIALOGUE then uses receptor-ligand interactions to propose cell-cell interactions that may mediate the novel multicellular programs that it identified (starting from a graph that depicts all the possible ligand-receptor interactions, and parsing it for each MCP to include its genes and short paths that link these genes; **Methods**). It further links the multicellular programs to genetic risk factors and to specific phenotypes of interest, thus providing a new way to uncover multicellular processes that mediate the genotype-to-phenotype mapping. As we show below, the multicellular programs identified by DIALOGUE are associated with complex phenotypes, ranging from animal behavior to clinical and pre-clinical disease manifestation.

### DIALOGUE identifies generalizable multicellular configurations and recovers spatial information from MERFISH in the mouse brain

To test DIALOGUE, we first applied it to MERFISH data, where the *in situ* expression of 155 genes was measured at the single-cell level in ∼1.1 million cells from the mouse hypothalamic preoptic region (10). We considered six broad cell types that were sufficiently represented in the data – microglia, endothelial cells, astrocytes, oligodendrocytes, excitatory and inhibitory neurons – and analyzed all pairwise combinations (**Supplementary Table 1**).

DIALOGUE identified corresponding transcriptional programs that were highly correlated across different cell types within the same physical niche (**Fig. 2A,B, right, light blue bars**). In contrast, single genes show only a moderate correlation across different cell types within the same niche (**Fig. 2C, Supp. Fig. 1A**), and standard dimensionality reduction approaches, such as principle component analysis (PCA) and Non-negative Matrix Factorization (NMF) did not reveal spatial associations across cell types (**Fig. 2C, Supp. Fig. 1A**).

**Figure 2.**
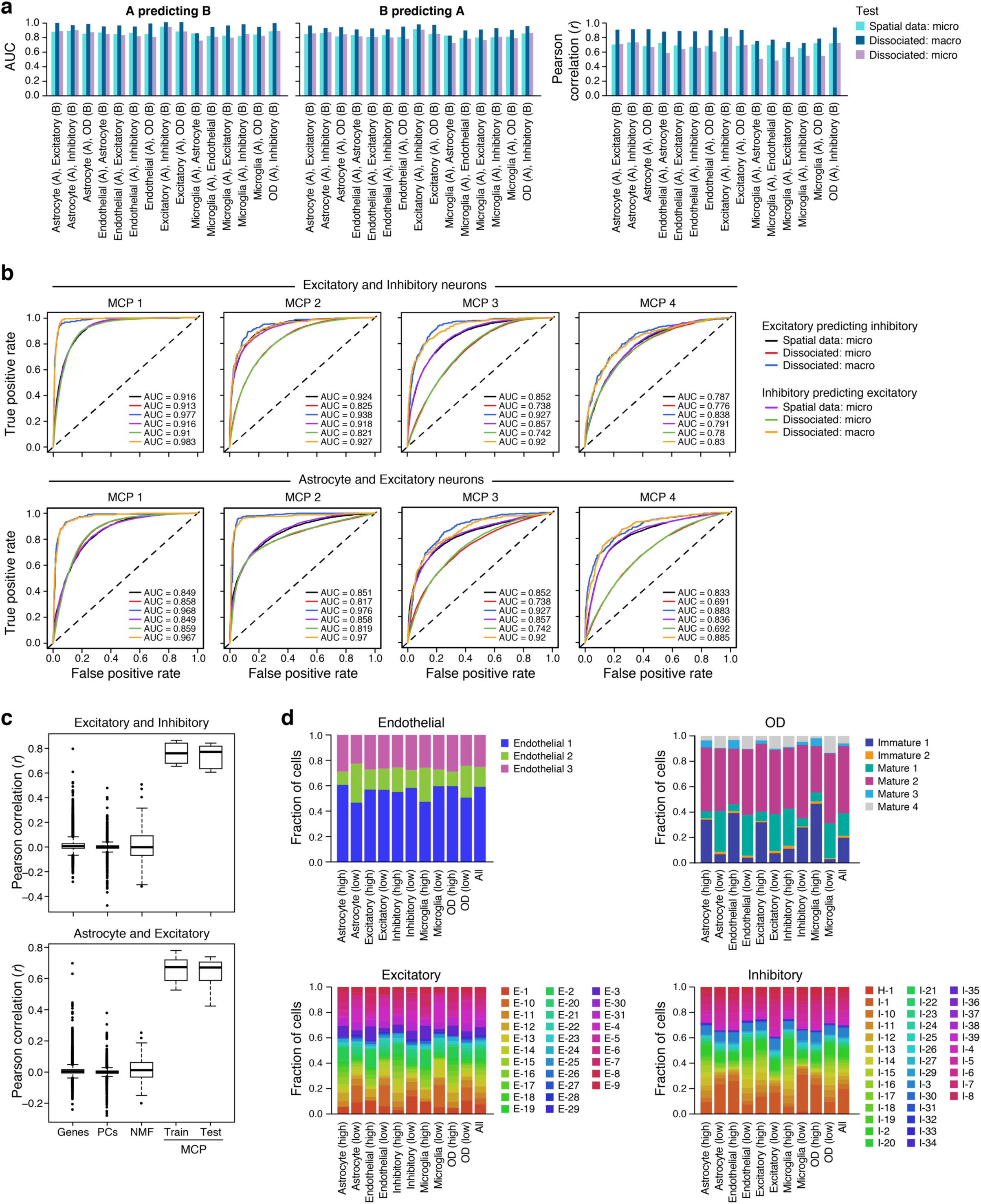
DIALOGUE identified multicellular programs across six cell types in the mouse hypothalamus and recovered spatial information at different scales. **(A,B)** Identification of MCPs at the micro and macro scales. **(A)** Pearson correlation coefficient (y axis, right) between the corresponding cell type programs, and performance (y axis, left and middle) when predicting the expression of the corresponding DIALOGUE component in the neighboring cells located in the same macro-environment (dark blue, ∼500 cells) or micro-environment (purple, and light blue, 15 cells), when testing on unseen test set; the training data includes either spatial coordinates and single-cell transcriptomes (light blue, “spatial data”) or only transcriptomes of ∼500 cell aggregates, without spatial information (“dissociated”, **Methods**). **(B)** The ROCs for predictions of the expression of each DIALOGUE program in unseen cells from the state of their neighbors in the “micro-” (15 cells) or “macro-” (∼500 cells) environment, when using full spatial coordinates in the training data (black and purple) or only the single-cell transcriptomes grouped to samples (red/blue, green/orange). **(C)** DIALOGUE recovers correlated programs across cell types. Pearson correlation coefficient between genes, PCs, NMF, or DIALOGUE MCPs from either the training or the test set (x axis) across different pairs of cell types in spatial niches in the mouse hypothalamus (see **Supp. Fig. 1A** for all other pairwise combinations). Middle line: median; box edges: 25^th^ and 75^th^ percentiles, whiskers: most extreme points that do not exceed ±IQR*1.5; further outliers are marked individually. **(D)** MCPs are not merely driven by cell subtype composition in a niche. The cells that over- or under-express the different MCP programs (top or bottom 25%, respectively, x axis) are composed of cells from different clusters (as previously defined (10), y axis); shown for the different cell types and their MCPs (i.e., MCP1).

The MCPs that DIALOGUE learned generalized well and had predictive value in a train-test setting. Specifically, we learned MCPs, each with two cell-type-specific components, from MERFISH data from a training set of 9 animals, and then tested whether these MCPs can predict the properties of the cell’s environment only from its own transcriptional state in a test set of 9 other animals. Indeed, using the learned model from the training data and the transcriptional state of a given cell (*e.g.*, excitatory neurons) in the test data, DIALOGUE accurately predicted the expression level of the corresponding cellular programs in the unseen neighboring cells (*e.g.*, microglia, astrocytes, etc., Pearson’s *r* > 0.69, P < 1*10^−30^; AUROC: 0.79 - 0.9, 0.82 median, **Fig. 2A,B**). We obtained similar results when withholding only males or only females (data not shown).

Next, we challenged DIALOGUE to recover such information by training it without any spatial coordinates, providing as input only samples that consisted of hundreds of cells from one tissue “patch” (“macro-environments” ∼500 “*in silico* dissociated” cells; ∼50-100µm^2^ each, **Methods**). We applied it on this cohort as before to learn programs for all pairs of cell types. Despite not having spatial data for training, DIALOGUE nonetheless was able to use the cell’s expression to predict the expression of the multicellular program in its unseen neighbors when tested on data from 9 other animals, which were not used in any part of the analysis other than to evaluate these predictions. For example, it successfully used the expression of inhibitory neurons to predict the state of their neighboring astrocytes (**Fig. 2B**). DIALOGUE performed well in this task both when predicting the microenvironment features (direct neighbors in a radius of ∼15 cells; AUROC: 0.73 - 0.91, median 0.82; P < 1*10^−30^, Mann-Whitney test) and macroenvironment features (AUROC: 0.83 - 0.98, median 0.94; P < 1*10^−30^, Mann-Whitney test, **Fig. 2A,B**).

Importantly, DIALOGUE MCPs do not simply mirror cell sub-clusters, nor do they only reflect simple compositional variation across space, where different cell subtypes are localized in different regions in the hypothalamus. Specifically, when considering cell clustering based on previous analyses of this cohort (10) we find that cells with a high, low, or moderate expression of the different MCPs, span multiple clusters, and can be found as over- or under-expressed in each of these clusters (**Fig. 2D**). Second, the co-expression of the programs’ components remained strong even when considering only pairs of specific cell subtypes (*e.g.*, inhibitory neurons I-1 and excitatory neurons E-2; **Supp. Fig. 1B**) or after regressing out cell-subtype-specific variation in each cell type (**Supp. Fig. 2**). The MCPs thus reflect cellular programs that are distinct from those identified using standard dimensionality reduction and clustering procedures (10), and highlight variation in specific cellular components as opposed to only gross changes in tissue composition at the cell subtype level.

### The multicellular programs in the mouse hypothalamus are linked to animal behavior

Many of the MCPs that DIALOGUE identified were strongly associated with particular animal behaviors. This was observed mostly for the leading MCP (MCP1; P < 0.05 for 12 out of 15 first MCPs, mixed-effects, BH FDR (37), **Methods**; **Fig. 3A**), which we focused on here (**Fig. 3B,C**). Subsequent MCPs (MCP2-5) showed higher intra-animal variation, resulting from spatial co-variation within the tissue (**Fig. 3D,E**).

**Figure 3.**
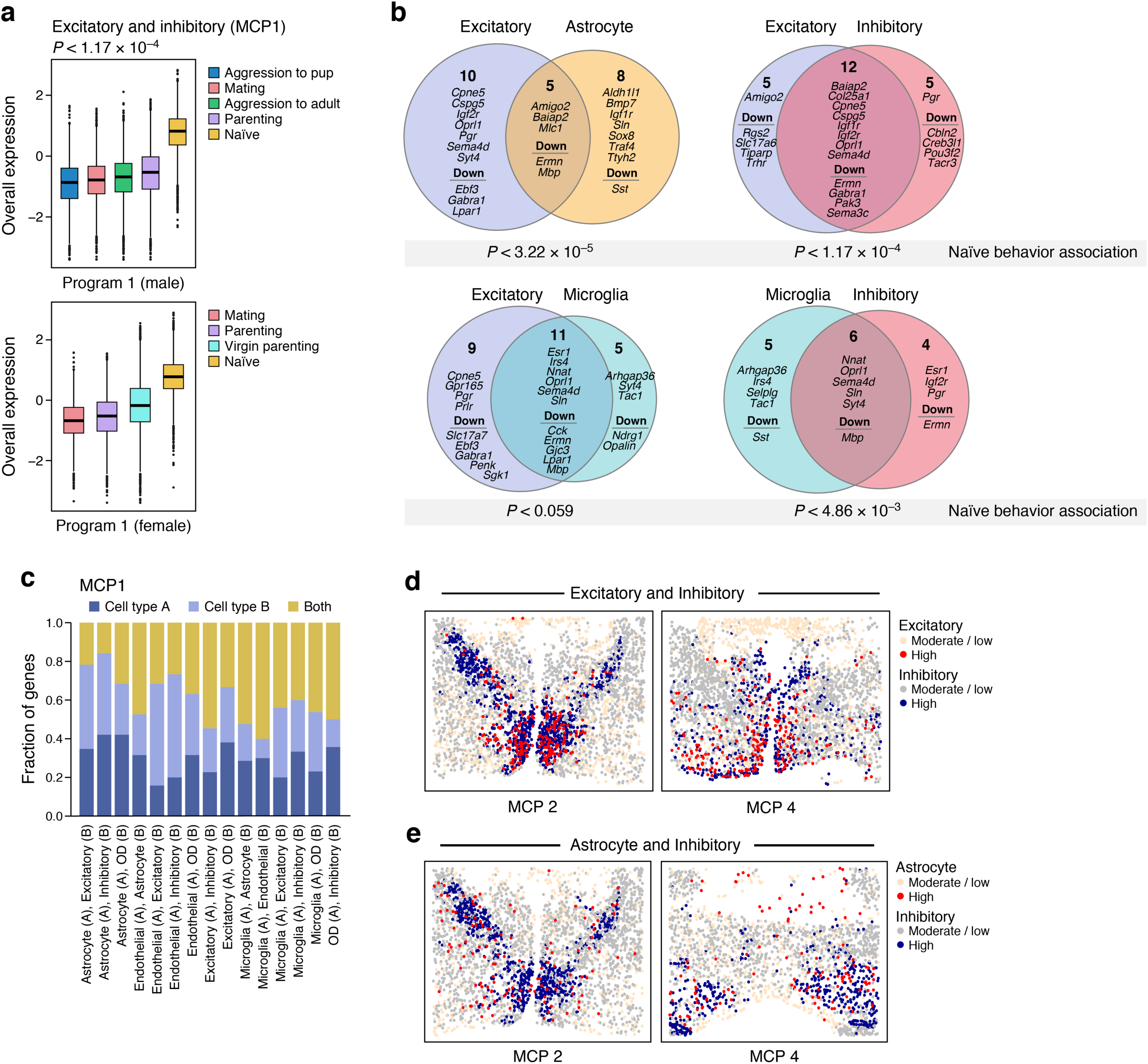
Shared and cell type specific components in DIALOGUE multicellular programs in the mouse hypothalamus. **(A)** Association of an excitatory-inhibitory program with animal behavior. Overall Expression (y axis, **Methods**) of the excitatory-inhibitory MCP in male (top) or female (bottom) mice exhibiting different behaviors (y axis). Middle line: median; box edges: 25^th^ and 75^th^ percentiles, whiskers: most extreme points that do not exceed ±IQR*1.5; further outliers are marked individually. P-values: multilevel mixed-effects models (**Methods**). **(B,C)** Shared and cell type specific genes in hypothalamus MCPs (shown for MCP1). **(B)** For each example hypothalamus MCPs, show are the two cell programs in the MCP and their specific and shared (intersection) genes. P-values: multilevel mixed-effects models. **(C)** Fraction of genes (y axis) that are shared (yellow) or specific to one (A, dark blue) or another (B, light blue) of the cell programs in each of the hypothalamus MCPs (x axis). **(D-E)** Spatial distribution of MCPs. Expression (color code) of MCP2 and MCP4 components in single cells (dots, x and y coordinates) identified for excitatory and inhibitory neurons (D) and astrocytes and inhibitory neurons (E).

The first MCPs (MCP1) that oligodendrocytes, excitatory and inhibitory neurons formed with other cell types were strongly repressed after parenting, aggression, or mating (P < 0.05, mixed-effects, BH FDR (37), **Fig. 3A,B**). All of these programs include *Oprl1*, encoding the nociceptin receptor, aligned with the key role of nociceptin in learning and emotional behaviors (38). We also find the involvement of hormonal signals, where the microglia-excitatory neurons MCP includes the genes encoding the receptors for prolactin (*Prlr*, neurons), progesterone (*Pgr*, neurons) and estrogen (*Esr1*, both compartments; **Fig. 3B**), with similar patterns in the microglia-inhibitory neuron MCP (**Fig. 3B**). These “hormonal” programs were equally active in female and male mice (P > 0.1, mixed-effects) and their neuronal and microglia components were strongly co-regulated even when considering only male or only female animals (*r* > 0.63, P < 1*10^−30^, Pearson correlation).

In contrast to the MCPs DIALOGUE identified, spatially-independent components did not show any significant association with animal behavior. To recover these spatially-independent components of cell states, we regressed out the spatially-associated programs from the gene expression and computed the PCs of the residuals (**Methods**). None of these residual components were associated with animal behavior, indicating that a large fraction of the biologically meaningful cell-cell variation in this system is captured by the MCPs identified.

Further examining the genes comprising the MCPs (**Supplementary Table 1**), we see that different programs have unique features that may reflect specific intercellular cross-regulation, while some genes recur across MCPs, either exclusively within the same cell type but in different MCPs, or across different types (**Fig. 3C**). For example, *Selplg* was in the microglia compartment in 4 out of the 5 (pairwise) MCP1s that involved microglia, but was not found in other cell types. *Selplg* encodes P-selectin glycoprotein ligand-1 (SELPLG), that, together with P-Selectin, constitutes a receptor/ligand complex involved in the recruitment of activated lymphocytes, and it is one the topmost (ranked 4) repressed genes in disease associated (DAMs) *vs*. homeostatic microglia (39). Other genes, such as *Sox8* and *Ttyh2*, were repeatedly and specifically members of the astrocyte component of multiple MCPs. In contrast, some genes, as *Mbp* (myelin basic protein (40)) and *Nnat* (involved in brain development and synaptic plasticity (41)), were found in multiple cell types across *multiple* MCPs (**Fig. 3B**), suggesting that their expression may be spatially regulated in a cell-type-independent manner.

### DIALOGUE recovers MCPs across five cells types from scRNA-seq data from colon biopsies of healthy individuals and ulcerative colitis patients

Next, we applied DIALOGUE to scRNA-Seq data that was collected from dissociated tissue specimens, without cell-level spatial coordinates, treating each sample as one “niche”. The dataset (16) consisted of 366,650 single-cell RNA-Seq profiles from 68 colonoscopic biopsies (each ∼2.4 mm^2^) from 12 healthy individuals and 18 UC patients. At least two biopsies were obtained from each individual, such that in UC patients, one inflamed or ulcerated tissue (‘‘inflamed’’) and one of adjacent normal tissue (‘‘non-inflamed’’). In addition, each sample was further separated to ‘‘epithelial’’ (EPI) and ‘‘lamina propria’’ (LP) fractions, resulting in 115 spatially distinct samples across 30 individuals. We focused on five well-represented cell subsets – macrophages, transit amplifying (TA) intestinal epithelial cells (TA1 and TA2), CD8^+^ T cells and CD4^+^ T cells. Going beyond pairwise associations, we applied DIALOGUE in multi-way mode to identify co-regulated multicellular programs that span all five cell types simultaneously (**Supplementary Table 2, Methods**).

To test DIALOGUE in this setting, we first examined if it can identify “mis-localized cells” based on the assumption that such cells will not fit the state predicted by their macroenvironment, as reflected by DIALOGUE’s multicellular programs. We trained DIALOGUE with an “in silico contaminated” dataset we generated, where we added to each sample ∼50 cells (10 per cell type; 0.02-0.06% of the cell type) from an “adjacent sample” from the same individual. These “adjacent samples” could be from the same or different layer (LP/EPI), and have the same or a different clinical status (both healthy, or inflamed and uninflamed). Based on the resulting multicellular programs that DIALOGUE found in the “contaminated training set”, it then computed an *environment-score*, which denotes for each cell to what extent its real state agrees with the one predicted by its neighbors (some of which may be contaminating; **Methods**).

DIALOGUE identified the mis-localized cells with high accuracy (**Fig. 4A**), as their environment-scores were significantly lower than those of the other cells (P < 1*10^−10^, *t*-test). We further evaluated this when considering different types of contamination, and found that DIALOGUE was most accurate in spotting contamination from a different and distinct spatial location (*i.e.*, LP *vs*. EPI, **Supp. Fig. 3**), and most successful for CD8 and CD4 T cells, and least so for macrophages (**Fig. 4A, Supp. Fig. 3**). These findings demonstrate the robustness of the approach to data contamination and the potential of using it to identify “infiltrators” to an established environment.

**Figure 4.**
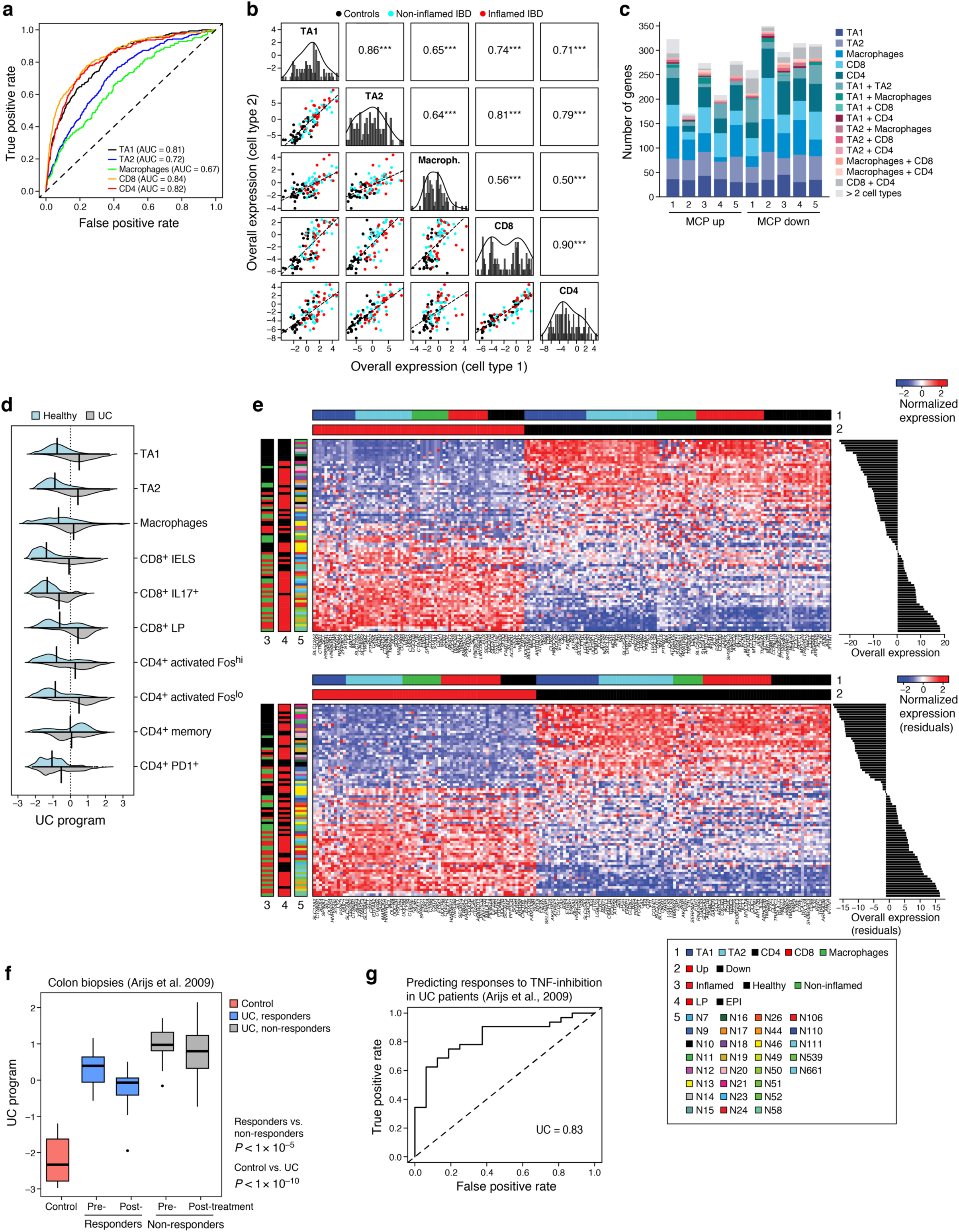
DIALOGUE multicellular programs associated with ulcerative colitis and predictive of clinical outcomes. **(A)** Deviation from multicellular patterns identifies mis-localized cells. ROC curve of the true positive (y axis) and false positive (x axis) rate when predicting mis-localized cells of each cell subset (color legend). **(B)** MCP1 in the human colon. Off diagonal panels: Comparison of Overall Expression scores (y and x axes) for each cell component of MCP1 (rows and columns, labels on diagonal) across the samples (dots, black: control, blue: non-inflamed IBD, red: inflamed IBD); the lines correspond to the linear fit. Pearson correlation (*r*) and significance (^***^P < 1*10^−3^) are shown in the panels above the diagonal. Diagonal panels: Distribution of Overall Expression scores for each cell type component, along with the kernel density estimates. **(C)** Most genes in five colon MCPs are specific to one cell component. Number of genes (y axis) in the up (left bars) and down (right bars) of each of five colon MCPs (x axis) that appear only in one cell type component or in multiple ones (color legend). **(D)** MCP1 is induced in UC samples. Distribution of Overall Expression scores (y axis) of MCP1 (“UC program”) across the different cell subtypes (*y* axis) for cells from UC patients (grey) and healthy individuals (light blue). **(E)** UC multicellular program genes. Top heatmap: Average expression (Z score, red/blue color bar) of top genes (columns) from the UC multicellular program, sorted by their pertaining cell type (top color bar), across samples (rows), sorted by Overall Expression (right, **Methods**), and labeled by clinical status, location and patient ID (left color bar). Program scores on bottom heatmap are after regressing out the impact of tissue location (LP/EPI), and selective the top genes based on these scores. **(F,G)** UC multicellular program predicts response to anti-TNF therapy. **(F)** Overall Expression (y axis) of the UC-program in bulk RNA-seq of colon biopsies from 24 UC patients pre- and post-infliximab infusion, stratified to responders (blue) and non-responders (grey) and in normal mucosa from 6 control patients (50) (red middle line: median; box edges: 25^th^ and 75^th^ percentiles, whiskers: most extreme points that do not exceed ±IQR*1.5; further outliers are marked individually. P-value: linear regression model (**Methods**). **(G)** ROC curve for using the UC multicellular program score in pre-treatment samples (50) to predict the subsequent clinical responses to infliximab infusion in UC patients.

Applying our approach to the real (non-contaminated) data we identified five MCPs, each with five cell-type-associated components. The different cell-type components of each multicellular program were strongly co-expressed across the samples (**Fig. 4B**; P < 1*10^−3^, mixed-effect model that accounts for gender, spatial compartment (EPI/LP), and cellular subtypes), even when considering only samples from UC patients or from healthy individuals (P < 1*10^−3^, mixed-effects). The majority of genes in each MCP were cell type specific (59-87%), although some were shared between the components of two or more cell types (**Fig. 4C, Supplementary Table 2**), potentially representing the context-dependent and -independent impact that the environment has on different cells, respectively.

### A multicellular program in ulcerative colitis enriched for genetic risk loci and predictive of clinical outcome

The first multicellular program (MCP1) that DIALOGUE identified in the colonoscopic biopsies was substantially higher in UC samples compared to samples from healthy individuals (P < 1*10^−12^, mixed-effects; **Fig. 4C-E, Supp. Fig. 4**). This was observed also when considering only the inflamed or non-inflamed UC samples (P < 1*10^−6^, mixed-effects), only specific cell types, as well as finer T cell subsets (**Fig. 4D**). This program, which we termed the UC program, was enriched with genes located in genetic risk loci for IBD and UC (P < 1*10^−5^ and 1*10^−3^, hypergeometric test, when including either 57 high-confidence or a larger set of 333 UC-associated genes (16), respectively; **Methods**), particularly in its up-regulated set, which included *FCGR2A* (42,43) (macrophages), *PRKCB* (CD8), *CCL20* (44) (TA1 and TA2), *SLC39A8* (TA2), *FOS* (TA1, TA2, and macrophages), *GPR65* (45) (CD4), and *ITLN1* (46) (TA1) – all have been shown to be strongly associated with UC. Of note, the majority of the genes in the UC program (>75%), including *PRKCB* (CD8) and *SLC39A8* (TA2), are cell-type specific in the program, although they are expressed in a wide variety of cells.

The UC multicellular program was further enriched for TNF response genes in its TA and CD4 compartments (P < 1*10^−8^, hypergeometric test; e.g., *EGR1/2/3, TNFAIP3*), and cytokines in its macrophage compartment (P < 1*10^−5^, hypergeometric test; *e.g., CCL18, CCL2, CXCL10, CXCL9*). Its CD8 and CD4 T cell compartments were enriched for genes previously associated with *CX3CR1*^+^ recently-activated effector memory or effector CD8 T cells and *TCF7*^+^ central memory CD4 T cells (47), respectively (P < 1*10^−8^, hypergeometric test); while significant, this was only a partial overlap (Jaccard index < 0.055). The downregulated part of the UC-program associated oxidative phosphorylation in TA1 and TA2 cells (P < 1*10^−12^, hypergeometric test) and exhaustion genes in CD8 T cells (P < 1*10^−8^, hypergeometric test).

In the ligand-receptor network of the UC multicellular program (**Supplementary Table 3**) we find direct (putative) interactions between *CD44* on TA1 cells and several of its ligands in macrophages (*MMP1, MMP9*, and *SPP1*) and TA2 cells (*FN1* and *HBEGF*). We also find 58 indirect interactions, where two program genes are interacting with the same ligand or receptor “mediator” gene. These mediators are enriched with UC risk genes (12 out of 333; P = 8.53*10^−3^, hypergeometric test), including *ITGAV* (48) and *ITGA4* (48,49).

Indicating its generalizability, the UC-multicellular program was associated with UC status also in an independent cohort of colon tissue samples obtained from 24 UC patients before and 4-6 weeks after their first treatment with a TNF-inhibitor (infliximab infusion) and in normal mucosa from 6 control patients (50). The UC multicellular program was substantially higher in the UC patients compared to the controls (P < 1*10^−10^, linear regression; AUC = 1) and was repressed by the treatment (P < 0.05, linear regression, **Fig. 4F**). Moreover, it was lower in the responders (P < 1*10^−5^, linear regression) and was predictive of responses (AUC = 0.83 and 0.88, when considering only the pre- or post-treatment samples, respectively, **Fig. 4G**). These findings demonstrate DIALOGUE’s ability to identify multicellular programs that characterize pathological cellular ecosystems and are linked to clinical phenotypes and subsequent clinical responses to interventions.

### A multicellular program associated with Alzheimer’s disease and the aging brain

Finally, we applied DIALOGUE to 80,660 single-nucleus RNA-seq (snRNA-seq) profiles from the prefrontal cortex of 48 individuals with varying degrees of Alzheimer’s disease (AD) pathology (17) (**Methods**). DIALOGUE identified a multicellular program that was substantially increased in AD patients (P < 1*10^−3^, mixed-effects, controlling for patient age and gender, **Fig. 5A-C, Supplementary Table 4**). The program spans components in inhibitory and excitatory neurons, oligodendrocytes, oligodendrocyte-precursor-cell (OPCs), astrocytes, and microglia, and its different cell type compartments are largely distinct, with minimal overlap (**Fig. 5D**; 90-95% of the MCP genes are in a single cell type component). Further supporting its connection to AD, its different components were substantially induced in the disease state also in an independent cohort of bulk brain tissue samples collected from 350 individuals, with definitive diagnosis of AD pathology or no AD pathology (51,52) (AUROC = 0.65, P = 1.47*10^−6^, regression model, accounting for individual age and gender, **Methods**).

**Figure 5.**
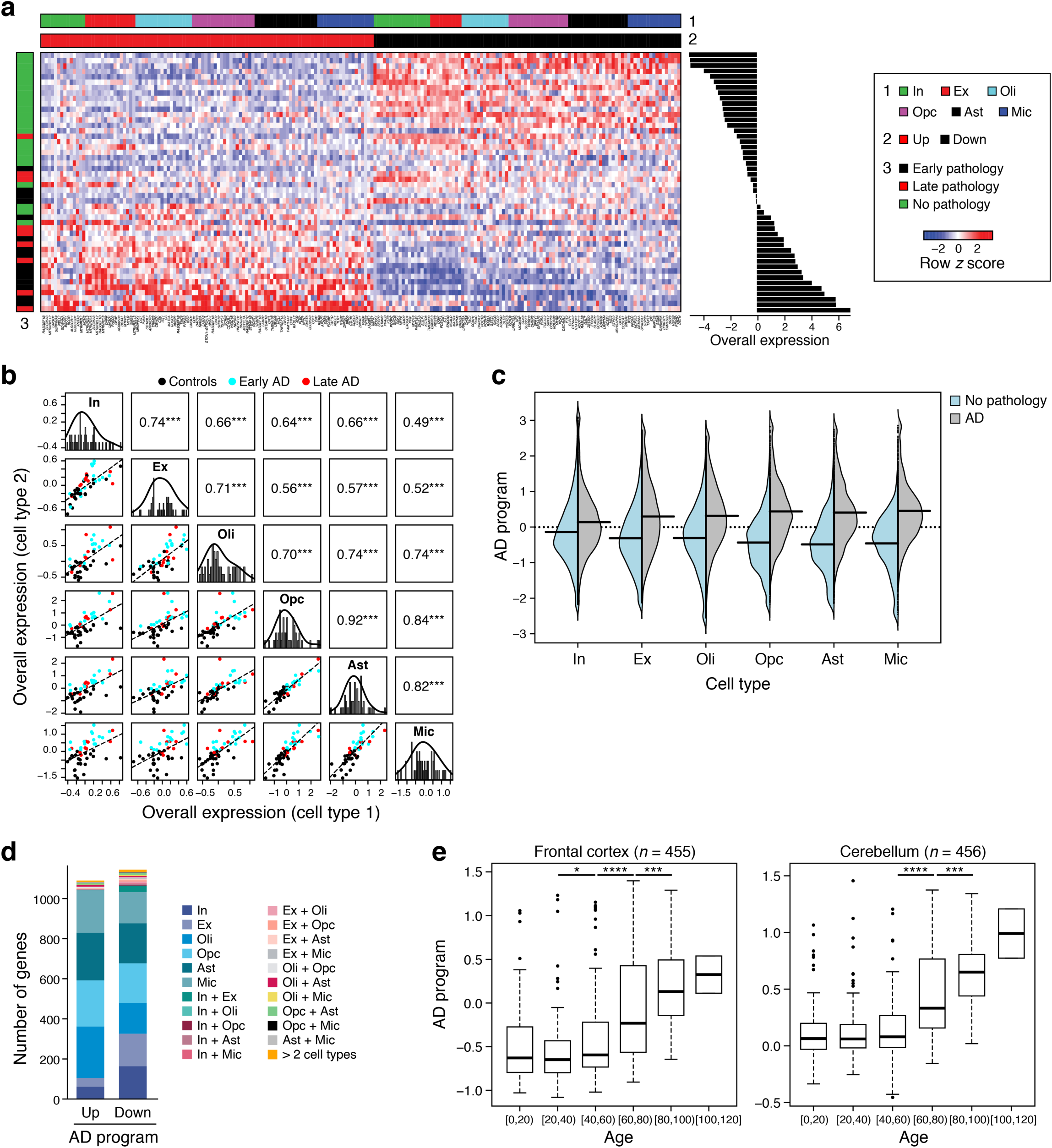
DIALOGUE identified a multicellular program in the prefrontal cortex that is associated with AD pathology and aging. **(A)** AD multicellular program. Average expression (Z score, red/blue color bar) of top genes (columns) from the AD multicellular program, sorted by their pertaining cell type (top color bar), across samples (rows), sorted by Overall Expression (right, **Methods**), and labeled by clinical status and patient ID (left color bar). **(B)** AD program components across cell types. Off diagonal panels: Comparison of Overall Expression scores (y and x axes) for each cell component of MCP1 (rows and columns, labels on diagonal) across the samples (dots, black: control, blue: early AD, red: late AD); the lines correspond to the linear fit. Pearson correlation coefficient (*r*) and significance (^***^P < 1*10^−3^) are shown in the upper triangle. Diagonal panels: Distribution of Overall Expression scores for each cell type component, along with the kernel density estimates. In: inhibitory neurons; Ex: excitatory neurons, Oli: oligodendrocytes, Opc: oligodendrocyte-precursor cells, Ast: astrocytes, Mic: microglia. **(C)** AD multicellular program components are induced across cell types in AD. Distribution of Overall Expression scores (*y* axis) of AD multicellular program component across the different cell subtypes (*y* axis) for cells from AD patients (grey) and neurologically normal subjects (light blue). **(D)** Most genes in the AD multicellular program are specific to one cell component. Number of genes (y axis) in the up (left bar) and down (right bar) compartment of the AD multicellular program (x axis) that belong to one cell type component or multiple ones (color legend). **(E)** AD multicellular program increases with age in the frontal cortex and cerebellum of neurologically normal subjects (see **Supp. Fig. 5**, where each of the program’s compartments is shown separately). Overall Expression of (y axis) of the AD multicellular program in bulk RNA-seq of the frontal cortex (left) and cerebellum (right) of neurologically normal subjects (61) stratified by age (*x* axis). Middle line: median; box edges: 25^th^ and 75^th^ percentiles, whiskers: most extreme points that do not exceed ±IQR*1.5; further outliers are marked individually. ^*^P<0.01, ^***^P<0.001, ^****^P<1*10^−4^ *t*-test.

This multicellular program includes the simultaneous repression of synaptic signaling genes in excitatory neurons (P < 1*10^−3^, hypergeometric test), of aerobic respiration genes in excitatory and inhibitory neurons (*e.g.*, the TCA cycle, P < 1*10^−5^, hypergeometric test), and of genes that facilitate proper synaptic activation in oligodendrocytes and astrocytes (*e.g.*, the excitatory amino acid transporter *SLC1A2*). Both its repressed and induced components include genes that have been previously linked to AD risk in genome-wide association studies (53–56) (13 and 14 genes, respectively; **Supplementary Table 4**), particularly in its astrocyte compartment (P < 0.05, hypergeometric test), including *APOE* (repressed in astrocytes and inhibitory neurons), *BZRAP1* (56) (astrocytes, repressed), HLA-DRA/B1 (57,58) (induced in astrocytes and repressed in excitatory neurons), *MEF2C* (59,60) (oligodendrocytes, repressed), and *MS4A4E* (oligodendrocytes, induced) (55).

As one of the greatest risk factors for Alzheimer’s is increasing age, we explored the possibility that the AD-program might become more prominent with age. Indeed, the up- and down-regulated parts of the program were also respectively enriched with genes that have been previously found to be over- or under-expressed in the aging cortex (P < 1*10^−5^, hypergeometric test). Moreover, the AD-program as a whole (**Fig. 5E**), and each of its cell-type compartments individually (**Supp. Fig. 5**), were strongly associated with age in bulk gene expression data that was obtained from both the frontal cortex (n = 455) and cerebellum (n = 456) of neurologically normal subjects across different ages (61). These findings thus demonstrate DIALOGUE’s ability to highlight pathological multicellular configurations that are tightly linked to different risk factors.

## DISCUSSION

In this study, we define the concept of multicellular programs and introduce it as a framework for studying tissue biology. We further develop the DIALOGUE method to recover multicellular programs, and apply it not only to spatial data but also to single cell genomics of dissociated tissue samples. The model it learns can predict the microenvironment of a cell – at different length scales – solely from its expression profile, and reveals multicellular programs associated to diverse phenotypes properties. DIALOGUE is entirely data-driven, and does not rely on strong underlying assumptions. It refers to different cell types as different representations or views of the same entity and maps multicellular programs directly from the data in an efficient and regularized way. The multicellular programs DIALOGUE identifies may arise due to shared (latent) factors in the cells’ micro- or macro-environment, due to shared (latent) genetic features, which impact different cell types in different ways, or due to a combination of genetic and environmental factors. All are instrumental for our understanding of homeostatic and disease states in tissue.

The AD- and UC-multicellular programs we identified included genes with germline variants associated with predisposition to these respective diseases, as well as genes that become increasingly over/under-expressed with age (in AD), drug treatment or response (in UC). Cell-intrinsic age-related processes leading to somatic genetic changes or epigenetic aberrations, together with the external cues from the aging microenvironment may impact different cell types in different yet correlated ways and give rise to such multicellular programs. Given additional information (*e.g.*, genotypes) or datasets where multiple samples are profiled from each individual, DIALOGUE can incorporate these sources of multicellular covariation and decouple their intertwined effects (**Methods**), discerning between genetic-related and environmental-related co-variation.

In the future, our approach can be extended to integrate the identified multicellular programs with models of transcriptional regulation (62) or intercellular signaling (30) to further explore their molecular basis. Integrating DIALOGUE with Perturb-Seq (63) data, especially when applied *in vivo* (64) (and measured *in situ*), or with human genetic data in larger cohorts, could provide a powerful approach to uncover the impact of genetic perturbations on the cell and its neighbors and set the stage for causal inference of gene function, vertically integrating from genetic causes, to single cells to tissue and clinical phenotypes.

While we focus here on multicellular transcriptional programs, DIALOGUE can be applied to any type of single cell data, from dissociated cells or *in situ*, including other molecular profiles or cell morphological features. Given multimodal data, it can identify both intracellular associations across different cellular modalities along with multicellular programs, and decouple the intracellular and multicellular processes that dictate different cellular properties (*e.g.*, cell shape, transcriptome, proteome, etc.).

DIALOGUE should be immediately applicable to existing datasets, allowing to extract multi-cellular programs from existing data, which was previously analyzed mostly at the single cell level. As the number of single-cell cohorts that are obtained from an increasing number of individuals and disease states grows rapidly, it should allow recovery of diverse cross-cell dependencies. More generally, the problem defined here should also prompt additional method development, and provide a basis to comprehensively map the vocabulary of multicellular programs underlying tissue function in health and disease.

## ACKNOWLEDGEMENTS

We thank Leslie Gaffney and Anna Hupalowska for help with figure preparation. L.J.A. is a CRI Irvington Fellow supported by the CRI, a fellow of the Eric and Wendy Schmidt Postdoctoral program and a recipient of the Burroughs Wellcome Fund CASI award. A.R. is an HHMI Investigator. Work was supported by the Klarman Cell Observatory, NIDDK RC2 DK114784, the Food Allergy Science Initiative, the Manton Foundation, and HHMI. The AD cohort was provided by the Rush Alzheimer’s Disease Center, Rush University Medical Center, Chicago. Data collection was supported through funding by NIA grants P30AG10161, R01AG15819, R01AG17917, R01AG30146, R01AG36836, U01AG32984, U01AG46152, U01AG61356, the Illinois Department of Public Health, and the Translational Genomics Research Institute.

## AUTHOR CONTRIBUTIONS

L.J. and A.R. conceived the study. L.J. devised the method and performed the analyses, with guidance and input from A.R. L.J. and A.R. wrote the manuscript.

## METHODS

### DIALOGUE. Overview

Given single-cell data obtained across different spatial locations or samples, DIALOGUE treats different types of cells from the same micro/macroenvironment or sample as different representations of the same entity (**Fig. 1**). It then identifies latent multicellular programs (MCPs), each composed of co-regulated gene sets that span multiple cell types, in two steps. In the first step, it uses PMD (36) to identify sparse canonical variates that transform the original feature space (*e.g.*, genes, PC, etc.) to a new feature space, where the different cell-type-specific representations are correlated across the different samples/environments. In the second step, given the new representation, DIALOGUE uses multilevel hierarchical modeling to identify the genes that comprise the latent features, while accounting for the single-cell distributions and controlling for potential confounders.

### DIALOGUE. Input

DIALOGUE takes as input single-cell profiles with either known spatial coordinates (from spatial methods) or with sample membership (*e.g.*, from a multi-sample scRNA-seq study). While we show applications for RNA profiles, other measurement type (e.g., protein levels, cell shape image-based features, etc.) can be used instead or in addition.

### DIALOGUE. PMD step

If the data is from dissociated tissue samples or biofluids, DIALOGUE first constructs for each cell type *z* a matrix *X*_*z*_, where (*X*_*z*_)_*ij*_ denotes the average value of feature *j* (*e.g.*, the expression of gene *j* or the value of PC *j*) in cell type *z* in sample *i*, following centering and scaling. Alternatively, if the data includes a cell’s spatial coordinates, a spatial niche is defined as a patch of a given number of cells, as few as two cells or as many as hundreds/thousands of cells. In this case, (*X*_*z*_)_*ij*_, denotes the average value of feature *j* in cells of type *z* in the spatial niche *i*. The number of features in each *X*_*z*_ can vary and does not have to be the same across the different cell types.

Given matrices *X*_1_, …, *X*_*N*_ representing *N* cell types, DIALOGUE applies PMD (36) to find sparse canonical variates *w*_1_, …, *w*_*N*_ by solving the following optimization problem

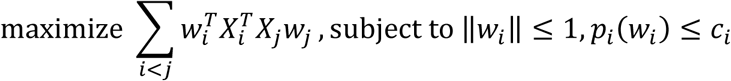

where *p*_*i*_ (*w*_*i*_) represent the LASSO penalties, and the tuning parameters *c*_*i*_ control the degree of sparsity. For each pair of cell types *i* and *j*, 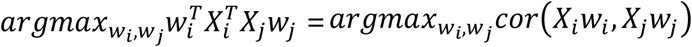, and therefore the resulting canonical variates identify a latent space where the new feature of cell type *i* (i.e., *X*_*i*_*w*_*i*_) are highly correlated with the new features of all the other cell types. The additional constrains ensure regularization and sparsity.

To select the tuning parameters, we defined 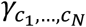 as the optimal value of the optimization function described above, when using a particular set of tuning parameters *c*_1_, …, *c*_*N*_. We defined 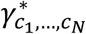 as the optimal value of the same optimization function and tuning parameters when using 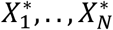 instead of the original dataset, where 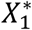 is the original data matrix *X*_*i*_ after permutation. For each set of candidate tuning parameters, we then permute the data multiple times and compute an empirical p-value based on the number of times that 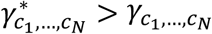. The tuning parameters that obtain the smallest p-value are selected.

In an iterative process termed multi-factor PMD (36), DIALOGUE identifies *K* latent features for each cell type, 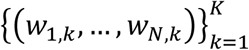. In practice, the new features of each cell type are orthogonal, minimizing redundancies, such that the greatest cross-cell-type correlations are observed when considering (*w*_1,1_, …, *w*_*N*,1_), gradually decreasing with increased *k*.

The new feature space is then defined at the single-cell level

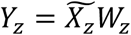

where 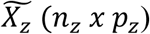 is the original feature space of cell type *z* (*e.g.*, normalized gene expression), and *W*_*z*_ (*p*_*z*_ *x K*) is the matrix of the sparse canonical variates identified for cell type *z*.

Of note, while the user can use different *K* values as input, DIALOGUE will always output the same *k*^th^ program for any *K* ≥ *k*.

### DIALOGUE. Multilevel modeling step

After DIALOGUE defined a new feature space, it identifies a gene signature for each sparse canonical variate, by interrogating the single-cell distributions, while accounting for potential confounding factors at different levels (*e.g.*, patient age, gender, sample type, cell sequencing quality, etc.).

For each of the *K* latent feature sets, DIALOGUE defines a set of *N* signatures (one per cell type), denoted as 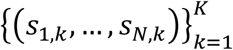.

A gene *g* is in *s*_*r,k*_ if its expression in cell type *r* is correlated with 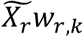 the pertaining latent feature of cell type *r*, and with 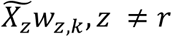, the pertaining latent features of the other cell types. The former is evaluated using partial Spearman correlation, when controlling for cell quality. The latter is tested using the following multilevel model

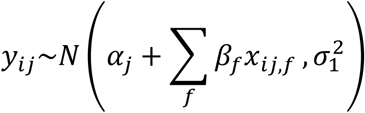

where *y*_*ij*_ is the value of the latent feature in cell *i* in sample *j, x*_*ij,f*_ is a cell-type-level covariate that controls for potential confounding factors (*e.g.*, log-transformed number of reads), and *α*_*j*_ is the sample *j* intercept, defined as

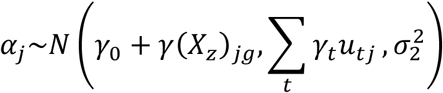

where *u*_*tj*_ are sample-level covariates of sample *j* (*e.g.*, patient age, gender), and (*X*_*z*_)_*jg*_ is the average expression of gene *g* in cell type *z* in sample *j*.

The model parameters and the pertaining p-values are computed using the lme4 (65) (https://cran.r-project.org/web/packages/lme4/index.html) and lmerTest R packages (66).

### DIALOGUE. Environment-score metric

The environment-score (*Es*) is defined based on the difference between the expected and observed cell state, given the identified multicellular programs.

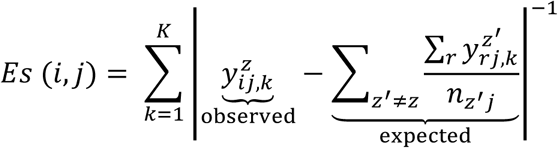

where 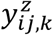 is the value of feature *k* in cell *i* of cell type *z* in sample *j*, and *n*_*z*′*j*_ is the number of cells of cell type *z’* in sample *j*.

To test if this measure can identify mis-localized cells, we used the colon/IBD data (16), where multiple samples were obtained from each individual. We performed an *in silico* contamination of the data by adding to each sample ∼50 cells (10 per cell type) from an “adjacent” sample obtained from the same individual. These “adjacent samples” could be from the same or different layer (LP/EPI), and have the same or a different clinical status (both healthy, or inflamed and uninflamed). Applied to this “contaminated” data, DIALOGUE identified MCPs spanning TA1, TA2, macrophages, CD8 and CD4 T cells, and computed the environment-scores as described above.

The predictions were tested per cell type (**Fig. 4A, Supp. Fig. 3**), when considering all cells, only control patients, only IBD patients, or only specific types of mis-localization, namely, erroneous tissue compartment (LP *vs*. EPI), same tissue type but different clinical status (considering only IBD samples, where inflamed and non-inflamed samples were collected for the same patient), and adjacent sample of the same tissue type (considering only control samples where similar samples were obtained from the same individual).

### DIALOGUE application to spatial transcriptomic data from the mouse hypothalamus

We applied DIALOGUE to single-cell transcriptomes of 155 genes across ∼1.1 million cells from the mouse hypothalamic preoptic region (10) measured by MERFISH. The processed data was downloaded from DRYAD (https://datadryad.org/stash/dataset/doi:10.5061/dryad.8t8s248), where expression values for the 135 genes measured in the combinatorial smFISH run were determined as the total counts per cell divided by the cell volume and scaled by 1,000. Expression values for the 21 genes measured in non-combinatorial, sequential FISH rounds were arbitrary fluorescence units per µm^3^, such that the same scale was used for all cells. The expression values were then centered and scaled for all genes. The gene *Fos* was not included in the analyses, as it was not measured in all cells, resulting in 155 genes.

The “behavior” annotation was used to examine the association of the programs with animal behavior. “Cell class” annotations were used to assign cells to major cell types (*e.g*., oligodendrocytes) and subtype (*e.g.*, immature oligodendrocytes 1), and the “neuron cluster_ID” was used to assign neurons to different neuronal subtypes (*e.g.*, I-1 for inhibitory neurons cluster 1, E-2 for excitatory neurons cluster 2, etc.).

Because the number of observations substantially exceeded the number of features in this cohort, we used the scaled and centered gene expression matrix as the “original features space” (**Fig. 1B**, “input”). This is equivalent to using all the Principle Components (PCs), as DIALOGUE is invariant to linear transformations. Niches were defined based on the two-dimensional coordinates of the cells’ centroid positions as patches of 15 or 500 adjacent cells, denoted as micro- and macro-environments respectively.

### DIALOGUE application to colon scRNA-Seq data

We applied DIALOGUE to scRNA-seq of 68 colonoscopic biopsies (each ∼2.4 mm^2^) from 12 healthy individuals and 18 UC patients (16). The processed UC data was downloaded from the single cell portal (https://singlecell.broadinstitute.org/single_cell/study/SCP259/intra-and-inter-cellular-rewiring-of-the-human-colon-during-ulcerative-colitis#study-download). Genes that were identified in the original study (16) as having a low signal to noise ratio due to putative ambient RNA contamination were removed. As the number of genes exceeded the number of samples in this cohort, the “original feature space” was the top 30 PCs, where PCs were computed based on the gene expression of the top 2,000 most variable genes, defined using the Seurat package *FindVariableFeatures* function. In this procedure, local polynomial regression (LOESS) is used to estimate the expected variance given the average gene expression values across the cells, on a log-log scale. Deviation from the expected value is then used to identify overdispersed genes.

### Definition of IBD risk genes

Two lists of IBD risk genes were used, as previously defined (16). The first list includes 333 genes that reside in region of linkage disequilibrium (LD) with IBD-associated-variants (44,48,67,68), after removing associations that map to more that 50 variants. The second list includes 57 putative risk genes with fine-mapped or nonsynonymous protein coding IBD or UC variant, and genes which were the only genes in the IBD-variant LD region.

### DIALOGUE application to AD snRNA-seq

DIALOGUE was applied to snRNA-seq data from the prefrontal cortex of 48 individuals with varying degrees of AD pathology (17). The data was previously generated as a part of the Religious Orders Study (ROS) or the Rush Memory and Aging Project (MAP) study (ROS/MAP). We downloaded the data from the AMP-AD Knowledge Portal (Synapse IDs: syn18686381, syn18686382, syn18686372) through controlled access, subject to the use conditions set by human privacy regulations.

To capture disease-related variation, which in this case was subtler and potentially masked by other sources of cell-cell variations, PCs were first computed based on genes that showed at least some moderate association with the disease state (t-test p-value < 0.1, without correction for multiple hypotheses). The top 30 PCs were used as the original feature space (**Fig. 1B**, “input”), ensuring that the number of features will not exceed the number of samples.

Bulk RNA-Seq data from the ROS/MAP study was used to examine the AD program in a larger cohort of 638 cortex autopsies (51,52). Data was downloaded from AMP-AD Knowledge Portal (Synapse IDs: syn3505720) through controlled access, subject to the use conditions set by human privacy regulations. For each sample the Overall Expression (OE) of the AD program was computed, using a scheme that filters technical variation and highlights biologically meaningful patterns, as described previously (18,69). Samples from patients with non-definitive diagnosis (CERAD score of 2 or 3) were discarded, and the association of the AD program OE with disease status (AD or non-AD, defined as CERAD scores of 1 and 4, respectively) was examined using a linear regression model that accounts for the individuals’ age and gender.

### Putative cell-cell interactions mediated by ligand-receptor binding

DIALOGUE starts from a graph of cognate ligand-receptor pairs (70). Given an MCP, each cell type is added to this ligand-receptor graph as another node, which is connected with edges to all of the genes in its compartment. Paths connecting the different cell types nodes may represent the molecular crosstalk that generated the multicellular co-expression pattern DIALOGUE identified. Paths of length three that connect different cell types represent direct ligand-receptor interactions, where the ligand is in the MCP compartment of one cell type and the receptor is in an MCP compartment of another cell type. Parsing the network to include only the cell type nodes and genes that are in short paths (< 5) between cell type nodes allows us to also examine potential mediators.

### Data availability

The datasets analyzed during the current study are available in the DRYAD repository https://datadryad.org/stash/dataset/doi:10.5061/dryad.8t8s248, single-cell portal https://singlecell.broadinstitute.org/single_cell/study/SCP259/intra-and-inter-cellular-rewiring-of-the-human-colon-during-ulcerative-colitis#study-download, and AMP-AD Knowledge Portal https://adknowledgeportal.synapse.org/ (Synapse IDs: syn18686381, syn18686382, syn18686372, and syn3505720; available through controlled access, subject to the use conditions set by human privacy regulations).

### Code availability

DIALOGUE is implemented as an R package and can be installed using the devtools::install (“DIALOGUE”) command. Further documentation and tutorials are provided in the package help pages (*e.g.*, ?DIALOGUE). We also provide DIALOGUE via GitHub (https://github.com/livnatje/DIALOGUE) and KCO repository (https://github.com/klarman-cell-observatory/DIALOGUE), along with additional guidelines and specifications.

**Supplementary Figure 1.**
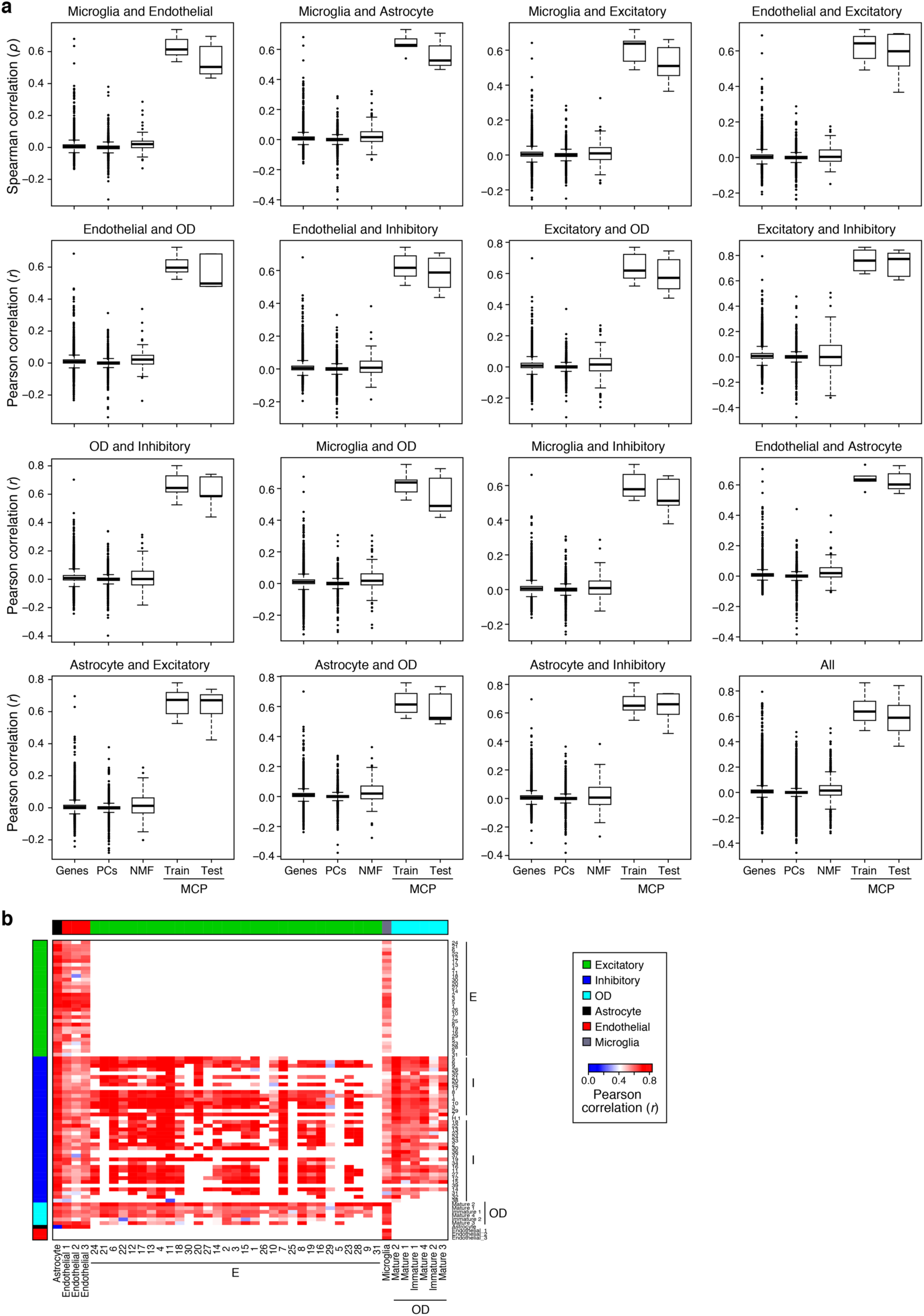
DIALOGUE identified multicellular programs in the mouse hypothalamus that are not recovered with other dimensionality reduction and clustering approaches. **(A)** Pearson correlation coefficient between genes, PCs, NMF, and DIALOGUE MCPs from either the training or the test set (*x* axis) across different pairs of cell types (panels) in spatial niches in the mouse hypothalamus. Middle line: median; box edges: 25^th^ and 75^th^ percentiles, whiskers: most extreme points that do not exceed ±IQR*1.5; further outliers are marked individually. **(B)** Pearson correlation coefficient (red/blue, color bar) between the Overall Expression of the relevant MCP component when considering only defined subsets of the pertaining cell types (rows, columns), as previously identified by clustering (10). White: missing values (cell subtypes that cannot be compared).

**Supplementary Figure 2.**
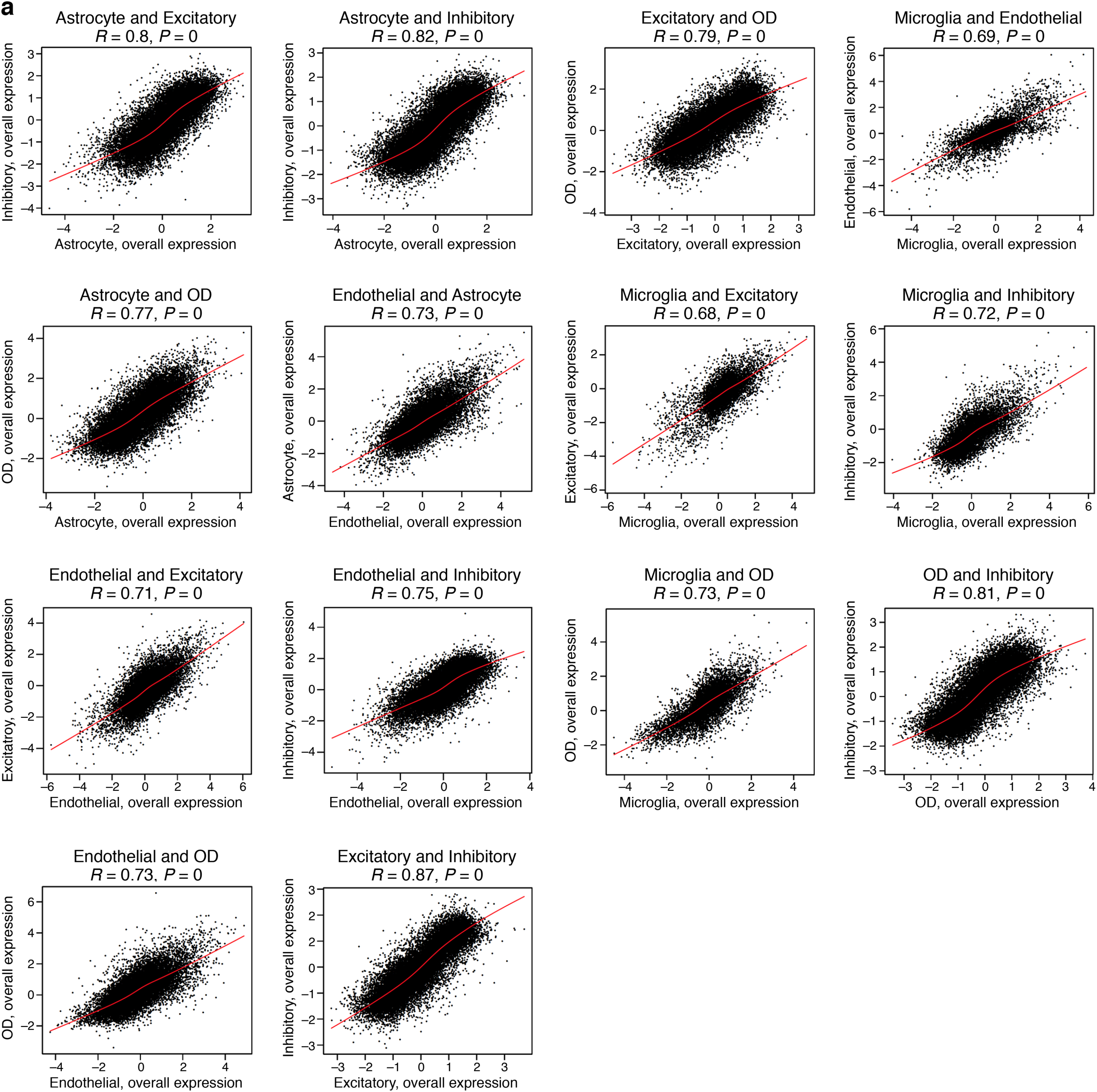
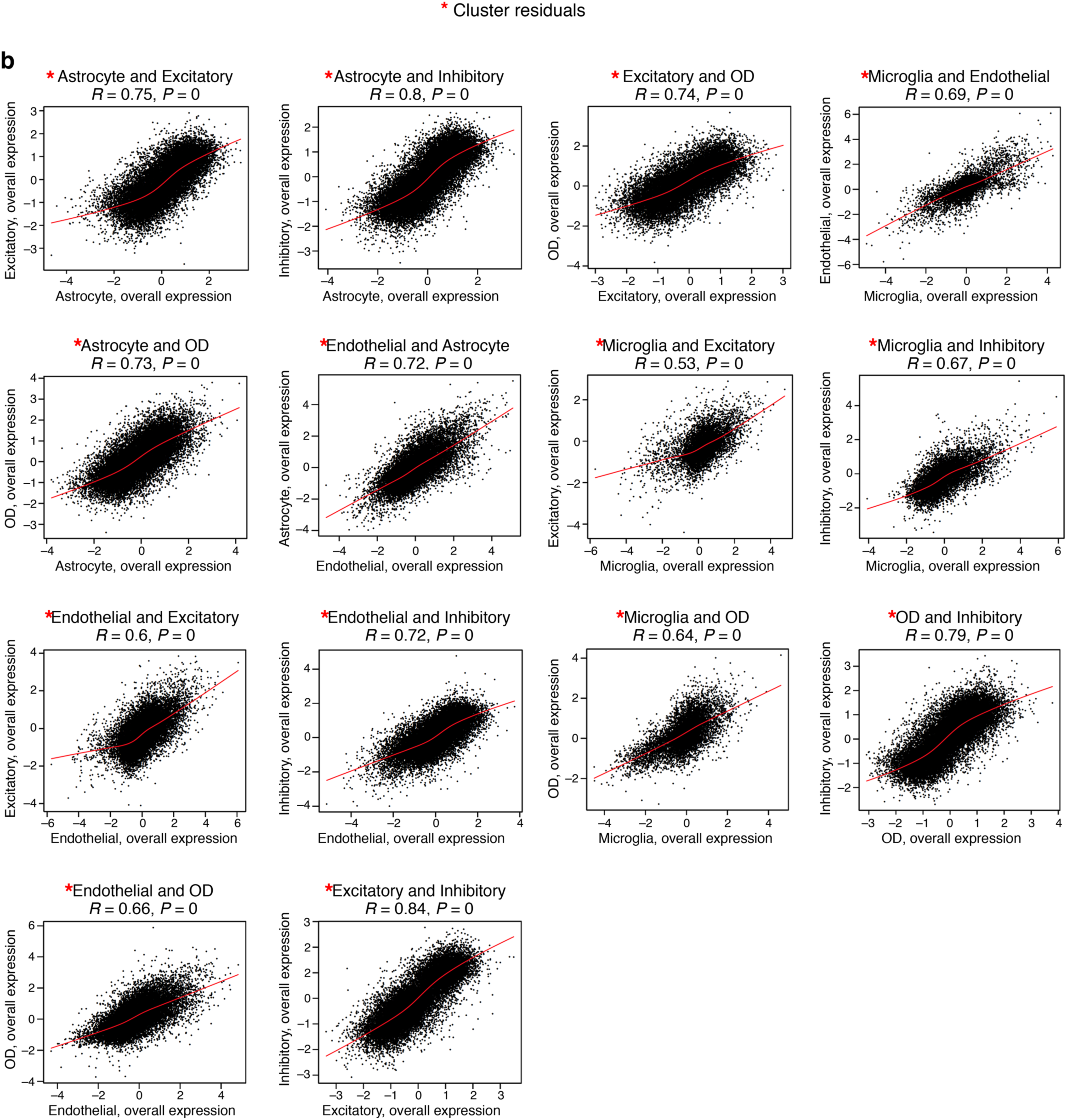
DIALOGUE’s MCPs are predictive of the cell’s environment. **(A)** Average Overall Expression in a niche (dot, 15 cells on average) of the first MCP (MCP1) in the first (x axis) and second (y axis) cell type in that MCP. **(B)** As in (A), but depicting the Overall Expression residuals after regressing out impact of cell clusters, as previously defined (10).

**Supplementary Figure 3.**
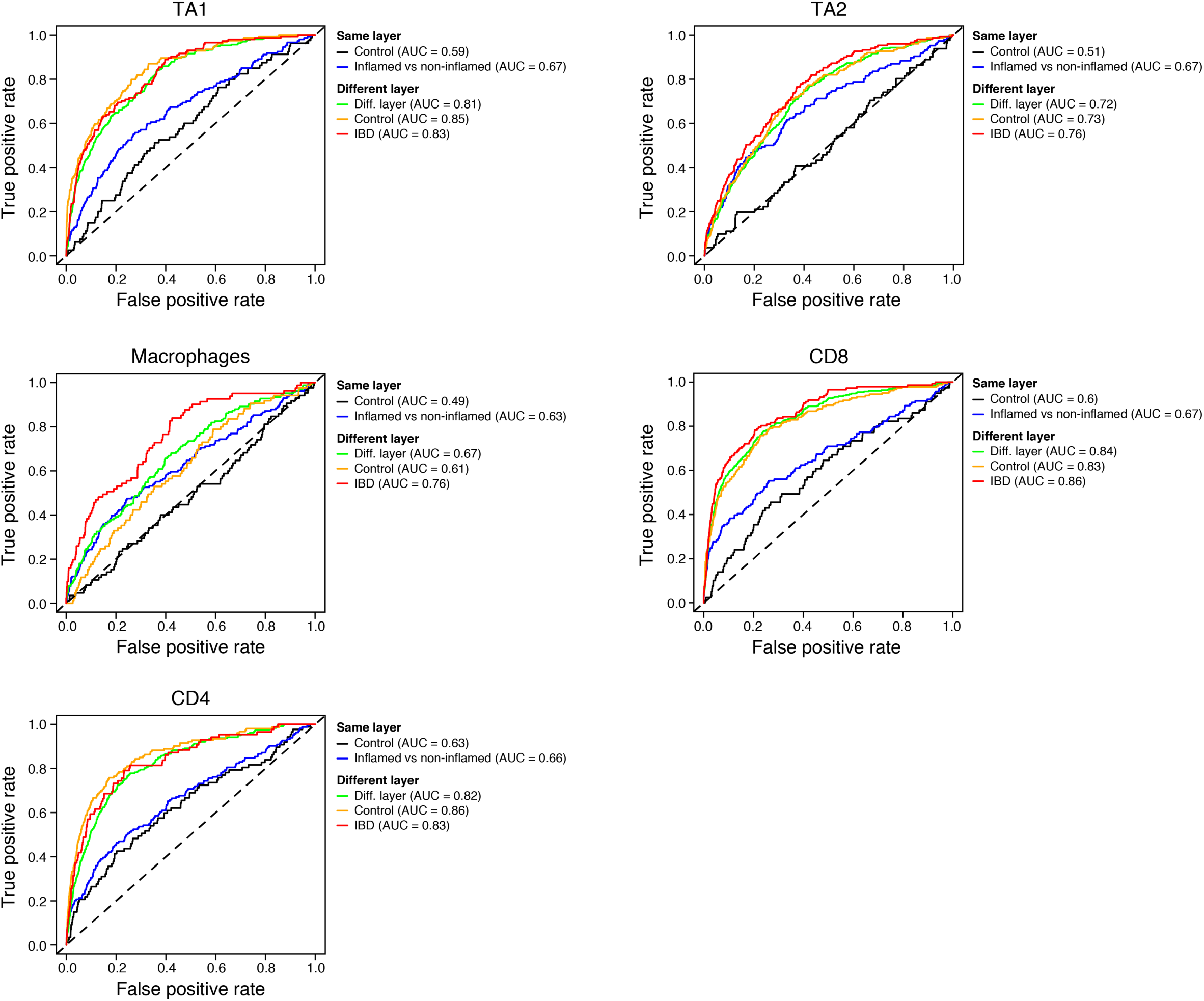
DIALOGUE’s ability to identify mis-localized cells varies by cell type and niche. ROC curve of the true positive (y axis) and false positive (x axis) rate when predicting mis-localized cells of each major subset (panels) with different types of “contamination” with cells that are either from the same layer (LP/EPI) within control (black, from replicate biopsy) or UC (blue; from adjacent biopsy with a different clinical status: inflamed or non-inflamed); or from a different layer but same clinical status, when considering either all samples (green) or only samples from control (yellow) or UC patients (red).

**Supplementary Figure 4.**
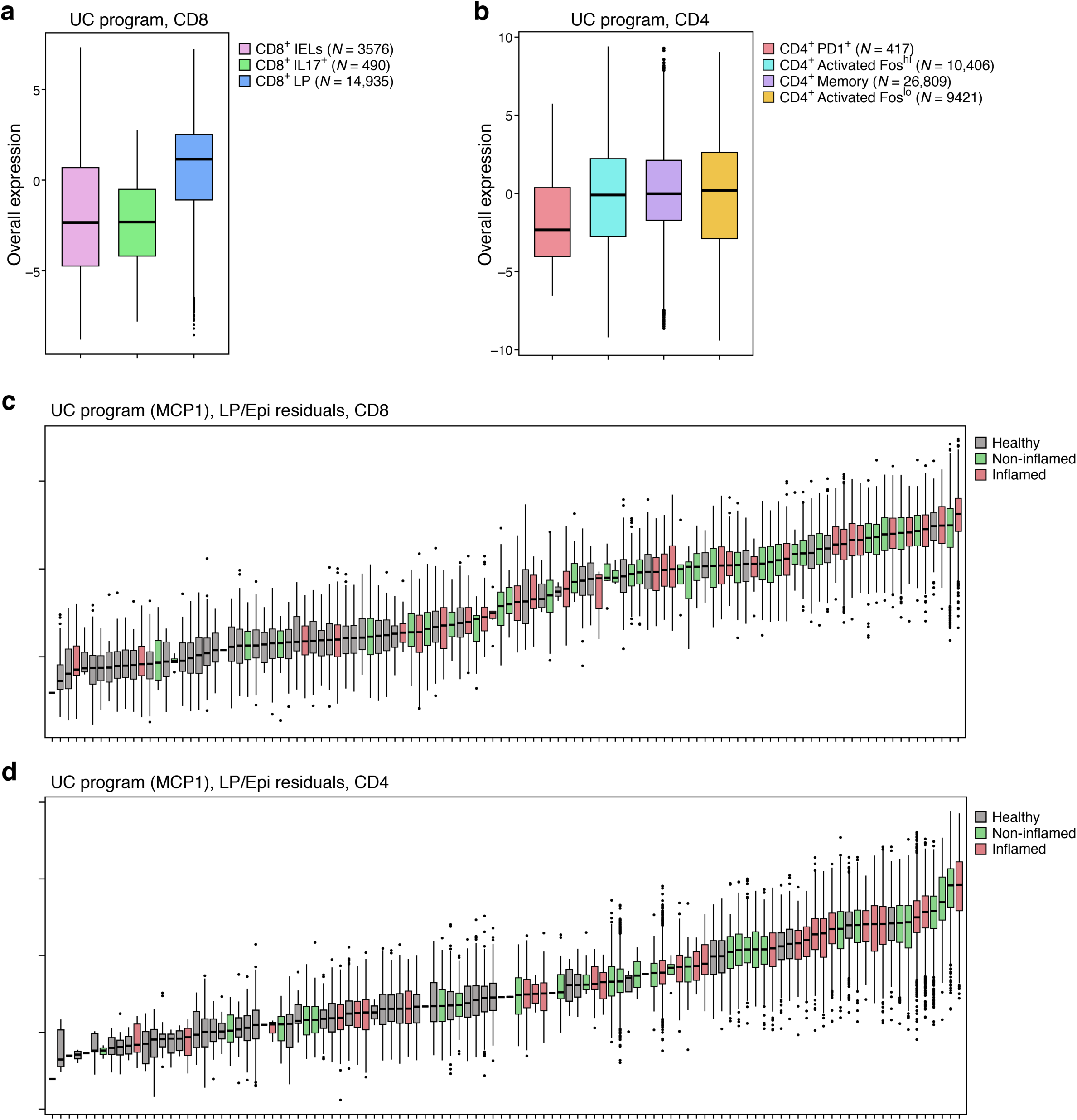
A multicellular program associated with ulcerative colitis. **(A,B)** Overall Expression (*y* axis) of the UC multicellular program component in CD8 (A) and CD4 (B) T cells, stratified by cell subtype (*x* axis). (**C,D**) Overall Expression (*y* axis) of the UC multicellular program in each sample (*x* axis) after regressing out the impact of the tissue layer (LP/EPI), in CD8 **(C)** and CD4 **(D)** T cells. Middle line: median; box edges: 25^th^ and 75^th^ percentiles, whiskers: most extreme points that do not exceed ±IQR*1.5; further outliers are marked individually.

**Supplementary Figure 5.**
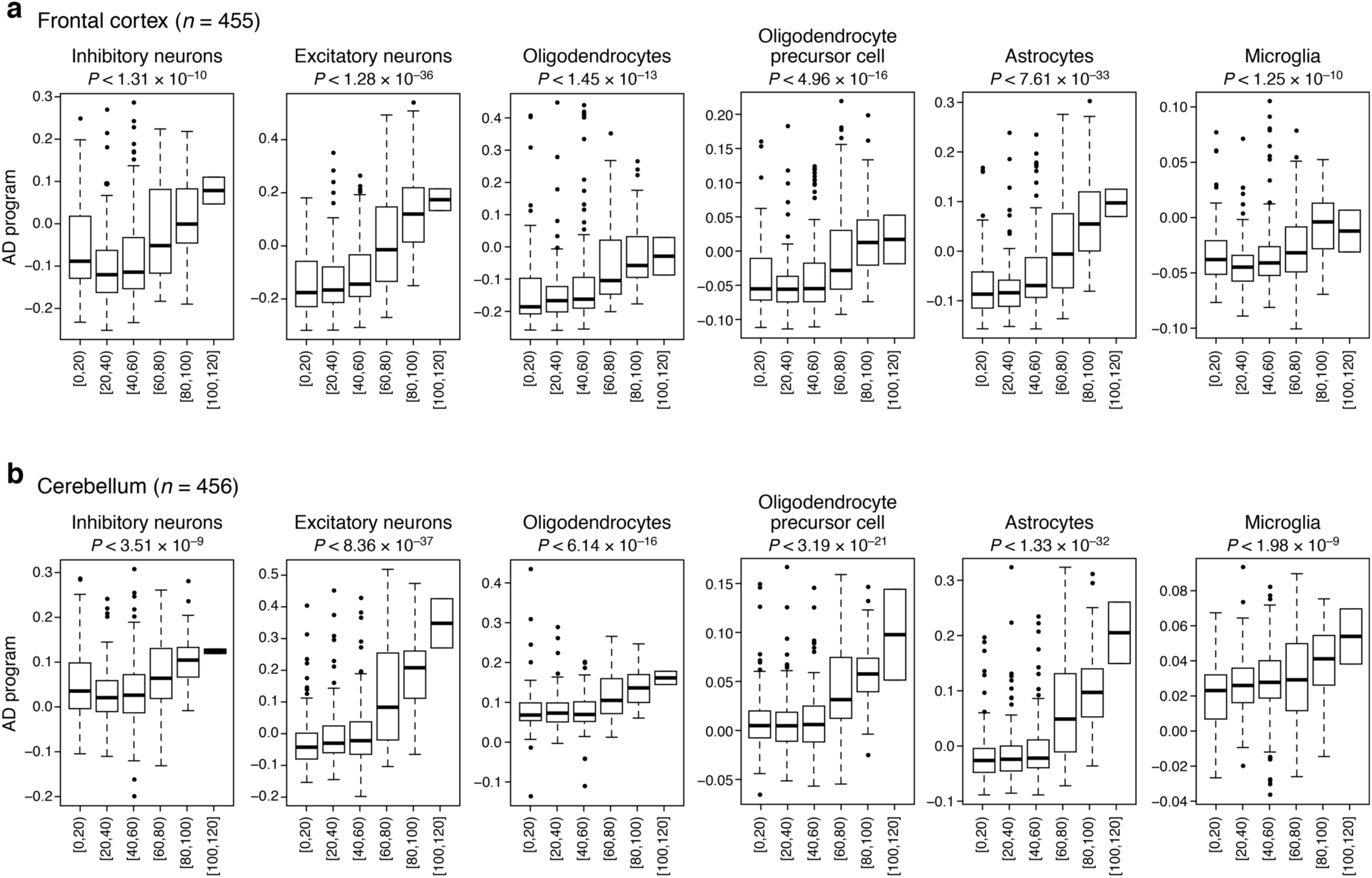
The different components of the AD multicellular program all mark brain aging. Overall Expression (*y* axis) of the different cell-type-specific AD multicellular program components in bulk RNA-seq of the frontal cortex **(A)** and cerebellum **(B)** obtained from neurologically normal subjects (61), stratified by age (*x* axis). Middle line: median; box edges: 25^th^ and 75^th^ percentiles, whiskers: most extreme points that do not exceed ±IQR*1.5; further outliers are marked individually. P-values computed based on a linear regression model (**Methods**).

## Supplementary Table Legend

**Supplementary Table 1.** MCPs identified in the mouse hypothalamus based on spatial transcriptomes (10).

**Supplementary Table 2.** MCPs identified in human colon based on scRNA-Seq data (16).

**Supplementary Table 3.** Ligand-receptor interactions associated with the UC program.

**Supplementary Table 4.** AD MCP identified based on single-nucleus data from brain autopsies (17).

## REFERENCES

1. Hong S, Stevens B. Microglia: Phagocytosing to Clear, Sculpt, and Eliminate. Dev Cell. 2016 Jul 25;38 (2):126–8.

2. Ribeiro M, Brigas HC, Temido-Ferreira M, Pousinha PA, Regen T, Santa C, et al. Meningeal γd T cell–derived IL-17 controls synaptic plasticity and short-term memory. Sci Immunol. 2019 Oct 11;4 (40):eaay5199.

3. Schwartz M. Can immunotherapy treat neurodegeneration? Science. 2017 Jul 21;357 (6348):254.

4. Baruch K, Deczkowska A, Rosenzweig N, Tsitsou-Kampeli A, Sharif AM, Matcovitch-Natan O, et al. PD-1 immune checkpoint blockade reduces pathology and improves memory in mouse models of Alzheimer’s disease. Nature Medicine. 2016 Jan 18;22:135.

5. Joyce JA, Fearon DT. T cell exclusion, immune privilege, and the tumor microenvironment. Science. 2015 Apr 3;348 (6230):74–80.

6. Corrigan-Curay J, Kiem H-P, Baltimore D, O’Reilly M, Brentjens RJ, Cooper L, et al. T-cell immunotherapy: looking forward. Mol Ther. 2014 Sep;22 (9):1564–74.

7. Hauser SL, Waubant E, Arnold DL, Vollmer T, Antel J, Fox RJ, et al. B-cell depletion with rituximab in relapsing-remitting multiple sclerosis. N Engl J Med. 2008 Feb 14;358 (7):676–88.

8. Macosko EZ, Basu A, Satija R, Nemesh J, Shekhar K, Goldman M, et al. Highly Parallel Genome-wide Expression Profiling of Individual Cells Using Nanoliter Droplets. Cell. 2015 May 21;161 (5):1202–14.

9. McDavid A, Finak G, Chattopadyay PK, Dominguez M, Lamoreaux L, Ma SS, et al. Data exploration, quality control and testing in single-cell qPCR-based gene expression experiments. Bioinformatics. 2013 Feb 15;29 (4):461–7.

10. Moffitt JR, Bambah-Mukku D, Eichhorn SW, Vaughn E, Shekhar K, Perez JD, et al. Molecular, spatial, and functional single-cell profiling of the hypothalamic preoptic region. Science. 2018 Nov 16;362 (6416):eaau5324.

11. Rodriques SG, Stickels RR, Goeva A, Martin CA, Murray E, Vanderburg CR, et al. Slide-seq: A scalable technology for measuring genome-wide expression at high spatial resolution. Science. 2019 Mar 29;363 (6434):1463.

12. Burgess DJ. Spatial transcriptomics coming of age. Nature Reviews Genetics. 2019 Jun 1;20 (6):317–317.

13. Kotliar D, Veres A, Nagy MA, Tabrizi S, Hodis E, Melton DA, et al. Identifying Gene Expression Programs of Cell-type Identity and Cellular Activity with Single-Cell RNA-Seq. bioRxiv. 2018 Jan 1;310599.

14. Zheng C, Zheng L, Yoo J-K, Guo H, Zhang Y, Guo X, et al. Landscape of Infiltrating T Cells in Liver Cancer Revealed by Single-Cell Sequencing. Cell. 2017 Jun 15;169 (7):1342-1356.e16.

15. Azizi E, Carr AJ, Plitas G, Cornish AE, Konopacki C, Prabhakaran S, et al. Single-Cell Map of Diverse Immune Phenotypes in the Breast Tumor Microenvironment. Cell. 2018 Aug 23;174 (5):1293-1308.e36.

16. Smillie CS, Biton M, Ordovas-Montanes J, Sullivan KM, Burgin G, Graham DB, et al. Intra- and Inter-cellular Rewiring of the Human Colon during Ulcerative Colitis. Cell. 2019 Jul 25;178 (3):714-730.e22.

17. Mathys H, Davila-Velderrain J, Peng Z, Gao F, Mohammadi S, Young JZ, et al. Single-cell transcriptomic analysis of Alzheimer’s disease. Nature. 2019 Jun 1;570 (7761):332–7.

18. Jerby-Arnon L, Shah P, Cuoco MS, Rodman C, Su M-J, Melms JC, et al. A Cancer Cell Program Promotes T Cell Exclusion and Resistance to Checkpoint Blockade. Cell. 2018 Nov 1;175 (4):984-997.e24.

19. Welch JD, Kozareva V, Ferreira A, Vanderburg C, Martin C, Macosko EZ. Single-Cell Multi-omic Integration Compares and Contrasts Features of Brain Cell Identity. Cell. 2019 Jun 13;177 (7):1873-1887.e17.

20. van der Maaten L, Hinton G. Visualizing Data using t-SNE. 2008;9 (Nov):2579--2605.

21. Fan J, Salathia N, Liu R, Kaeser GE, Yung YC, Herman JL, et al. Characterizing transcriptional heterogeneity through pathway and gene set overdispersion analysis. Nat Methods. 2016 Mar;13 (3):241–4.

22. Yang Z, Michailidis G. A non-negative matrix factorization method for detecting modules in heterogeneous omics multi-modal data. Bioinformatics. 2016 Jan 1;32 (1):1–8.

23. Vieth B, Parekh S, Ziegenhain C, Enard W, Hellmann I. A systematic evaluation of single cell RNA-seq analysis pipelines. Nature Communications. 2019 Oct 11;10 (1):4667.

24. Finak G, McDavid A, Yajima M, Deng J, Gersuk V, Shalek AK, et al. MAST: a flexible statistical framework for assessing transcriptional changes and characterizing heterogeneity in single-cell RNA sequencing data. Genome Biol. 2015 Dec 10;16:278–278.

25. Kharchenko PV, Silberstein L, Scadden DT. Bayesian approach to single-cell differential expression analysis. Nat Meth. 2014 Jul;11 (7):740–2.

26. Nitzan M, Karaiskos N, Friedman N, Rajewsky N. Gene expression cartography. Nature. 2019 Dec 1;576 (7785):132–7.

27. Cang Z, Nie Q. Inferring spatial and signaling relationships between cells from single cell transcriptomic data. Nature Communications. 2020 Apr 29;11 (1):2084.

28. Satija R, Farrell JA, Gennert D, Schier AF, Regev A. Spatial reconstruction of single-cell gene expression data. Nat Biotechnol. 2015 May;33 (5):495–502.

29. Achim K, Pettit J-B, Saraiva LR, Gavriouchkina D, Larsson T, Arendt D, et al. High-throughput spatial mapping of single-cell RNA-seq data to tissue of origin. Nat Biotechnol. 2015 May;33 (5):503–9.

30. Browaeys R, Saelens W, Saeys Y. NicheNet: modeling intercellular communication by linking ligands to target genes. Nature Methods. 2020 Feb 1;17 (2):159–62.

31. Kumar MP, D. J, Lagoudas G, Jiao Y, Sawyer A, Drummond DC, et al. Analysis of Single-Cell RNA-Seq Identifies Cell-Cell Communication Associated with Tumor Characteristics. Cell Rep. 2018 Nov 6;25 (6):1458-1468.e4.

32. Efremova M, Vento-Tormo M, Teichmann SA, Vento-Tormo R. CellPhoneDB: inferring cell–cell communication from combined expression of multi-subunit ligand–receptor complexes. Nature Protocols. 2020 Apr 1;15 (4):1484–506.

33. Edsgärd D, Johnsson P, Sandberg R. Identification of spatial expression trends in single-cell gene expression data. Nature Methods. 2018 May 1;15 (5):339–42.

34. Goltsev Y, Samusik N, Kennedy-Darling J, Bhate S, Hale M, Vazquez G, et al. Deep Profiling of Mouse Splenic Architecture with CODEX Multiplexed Imaging. Cell. 2018 Aug 9;174 (4):968-981.e15.

35. Haber AL, Biton M, Rogel N, Herbst RH, Shekhar K, Smillie C, et al. A single-cell survey of the small intestinal epithelium. Nature. 2017 Nov 1;551 (7680):333–9.

36. Witten DM, Tibshirani R, Hastie T. A penalized matrix decomposition, with applications to sparse principal components and canonical correlation analysis. Biostatistics. 2009 Jul;10 (3):515–34.

37. Benjamini Y, Hochberg Y. Controlling the False Discovery Rate: A Practical and Powerful Approach to Multiple Testing. Journal of the Royal Statistical Society Series B (Methodological). 1995;57 (1):289–300.

38. Andero R. Nociceptin and the nociceptin receptor in learning and memory. Prog Neuropsychopharmacol Biol Psychiatry. 2015 Oct 1;62:45–50.

39. Keren-Shaul H, Spinrad A, Weiner A, Matcovitch-Natan O, Dvir-Szternfeld R, Ulland TK, et al. A Unique Microglia Type Associated with Restricting Development of Alzheimer’s Disease. Cell. 2017 Jun 15;169 (7):1276-1290.e17.

40. Berger T, Rubner P, Schautzer F, Egg R, Ulmer H, Mayringer I, et al. Antimyelin antibodies as a predictor of clinically definite multiple sclerosis after a first demyelinating event. N Engl J Med. 2003 Jul 10;349 (2):139–45.

41. Joseph RM. Neuronatin gene: Imprinted and misfolded: Studies in Lafora disease, diabetes and cancer may implicate NNAT-aggregates as a common downstream participant in neuronal loss. Genomics. 2014 Feb 1;103 (2):183–8.

42. Asano K, Matsushita T, Umeno J, Hosono N, Takahashi A, Kawaguchi T, et al. A genome-wide association study identifies three new susceptibility loci for ulcerative colitis in the Japanese population. Nature Genetics. 2009 Dec 1;41 (12):1325–9.

43. Weersma RK, Crusius JBA, Roberts RL, Koeleman BPC, Palomino-Morales R, Wolfkamp S, et al. Association of FcgR2a, but not FcgR3a, with inflammatory bowel diseases across three Caucasian populations. Inflamm Bowel Dis. 2010 Dec;16 (12):2080–9.

44. Liu JZ, van Sommeren S, Huang H, Ng SC, Alberts R, Takahashi A, et al. Association analyses identify 38 susceptibility loci for inflammatory bowel disease and highlight shared genetic risk across populations. Nat Genet. 2015/07/20 ed. 2015 Sep;47 (9):979–86.

45. Tcymbarevich IV, Eloranta JJ, Rossel J-B, Obialo N, Spalinger M, Cosin-Roger J, et al. The impact of the rs8005161 polymorphism on G protein-coupled receptor GPR65 (TDAG8) pH-associated activation in intestinal inflammation. BMC Gastroenterol. 2019 Jan 7;19 (1):2–2.

46. Barrett JC, Hansoul S, Nicolae DL, Cho JH, Duerr RH, Rioux JD, et al. Genome-wide association defines more than 30 distinct susceptibility loci for Crohn’s disease. Nat Genet. 2008 Aug;40 (8):955–62.

47. Zhang L, Yu X, Zheng L, Zhang Y, Li Y, Fang Q, et al. Lineage tracking reveals dynamic relationships of T cells in colorectal cancer. Nature. 2018 Dec 1;564 (7735):268–72.

48. de Lange KM, Moutsianas L, Lee JC, Lamb CA, Luo Y, Kennedy NA, et al. Genome-wide association study implicates immune activation of multiple integrin genes in inflammatory bowel disease. Nat Genet. 2017/01/09 ed. 2017 Feb;49 (2):256–61.

49. Gerecke C, Scholtka B, Löwenstein Y, Fait I, Gottschalk U, Rogoll D, et al. Hypermethylation of ITGA4, TFPI2 and VIMENTIN promoters is increased in inflamed colon tissue: putative risk markers for colitis-associated cancer. J Cancer Res Clin Oncol. 2015 Dec;141 (12):2097–107.

50. Arijs I, Li K, Toedter G, Quintens R, Van Lommel L, Van Steen K, et al. Mucosal gene signatures to predict response to infliximab in patients with ulcerative colitis. Gut. 2009 Dec;58 (12):1612–9.

51. De Jager PL, Ma Y, McCabe C, Xu J, Vardarajan BN, Felsky D, et al. A multi-omic atlas of the human frontal cortex for aging and Alzheimer’s disease research. Sci Data. 2018 Aug 7;5:180142–180142.

52. Bennett DA, Buchman AS, Boyle PA, Barnes LL, Wilson RS, Schneider JA. Religious Orders Study and Rush Memory and Aging Project. J Alzheimers Dis. 2018;64 (1):S161–89.

53. Jansen IE, Savage JE, Watanabe K, Bryois J, Williams DM, Steinberg S, et al. Genome-wide meta-analysis identifies new loci and functional pathways influencing Alzheimer’s disease risk. Nature Genetics. 2019 Mar 1;51 (3):404–13.

54. da Rocha TJ, Silva Alves M, Guisso CC, de Andrade FM, Camozzato A, de Oliveira AA, et al. Association of GPX1 and GPX4 polymorphisms with episodic memory and Alzheimer’s disease. Neuroscience Letters. 2018 Feb 14;666:32–7.

55. Hollingworth P, Harold D, Sims R, Gerrish A, Lambert J-C, Carrasquillo MM, et al. Common variants at ABCA7, MS4A6A/MS4A4E, EPHA1, CD33 and CD2AP are associated with Alzheimer’s disease. Nat Genet. 2011 May;43 (5):429–35.

56. Tábuas-Pereira M, Santana I, Guerreiro R, Brás J. Alzheimer’s Disease Genetics: Review of Novel Loci Associated with Disease. Current Genetic Medicine Reports. 2020 Mar 1;8 (1):1–16.

57. Mansouri L, Klai S, Gritli N, Fekih-Mrissa N, Messalmani M, Bedoui I, et al. Association of HLA-DR/DQ Polymorphism With Alzheimer’s Disease. The American Journal of the Medical Sciences. 2015 Apr 1;349 (4):334–7.

58. Wang Z-X, Wang H-F, Tan L, Liu J, Wan Y, Sun F-R, et al. Effects of HLA-DRB1/DQB1 Genetic Variants on Neuroimaging in Healthy, Mild Cognitive Impairment, and Alzheimer’s Disease Cohorts. Molecular Neurobiology. 2017 Jul 1;54 (5):3181–8.

59. Boden KA, Barber IS, Clement N, Patel T, Guetta-Baranes T, Brookes KJ, et al. Methylation Profiling RIN3 and MEF2C Identifies Epigenetic Marks Associated with Sporadic Early Onset Alzheimer’s Disease. Journal of Alzheimer’s Disease Reports. 2017;1 (1):97–108.

60. Lambert JC, Ibrahim-Verbaas CA, Harold D, Naj AC, Sims R, Bellenguez C, et al. Meta-analysis of 74,046 individuals identifies 11 new susceptibility loci for Alzheimer’s disease. Nat Genet. 2013 Dec;45 (12):1452–8.

61. Hernandez DG, Nalls MA, Moore M, Chong S, Dillman A, Trabzuni D, et al. Integration of GWAS SNPs and tissue specific expression profiling reveal discrete eQTLs for human traits in blood and brain. Neurobiol Dis. 2012 Jul;47 (1):20–8.

62. Pratapa A, Jalihal AP, Law JN, Bharadwaj A, Murali TM. Benchmarking algorithms for gene regulatory network inference from single-cell transcriptomic data. Nature Methods. 2020 Feb 1;17 (2):147–54.

63. Dixit A, Parnas O, Li B, Chen J, Fulco CP, Jerby-Arnon L, et al. Perturb-Seq: Dissecting Molecular Circuits with Scalable Single-Cell RNA Profiling of Pooled Genetic Screens. Cell. 2016 Dec 15;167 (7):1853-1866.e17.

64. Jin X, Simmons SK, Guo AX, Shetty AS, Ko M, Nguyen L, et al. <em>In vivo</em> Perturb-Seq reveals neuronal and glial abnormalities associated with Autism risk genes. bioRxiv. 2019 Jan 1;791525.

65. Bates D, Mächler M, Bolker B, Walker S. Fitting Linear Mixed-Effects Models Using lme4. 2015th-10th–07 ed. Vol. 67, Journal of Statistical Software. 2015. p. 48.

66. Kuznetsova A, Brockhoff PB, Christensen RHB. lmerTest Package: Tests in Linear Mixed Effects Models. Journal of Statistical Software; Vol 1, Issue 13 (2017) [Internet]. 2017; Available from: https://www.jstatsoft.org/v082/i13

67. Huang H, Fang M, Jostins L, Umicevic Mirkov M, Boucher G, Anderson CA, et al. Fine-mapping inflammatory bowel disease loci to single-variant resolution. Nature. 2017 Jul 1;547 (7662):173–8.

68. Jostins L, Ripke S, Weersma RK, Duerr RH, McGovern DP, Hui KY, et al. Host– microbe interactions have shaped the genetic architecture of inflammatory bowel disease. Nature. 2012 Nov 1;491 (7422):119–24.

69. Jerby L, Neftel C, Shore ME, McBride MJ, Haas B, Izar B, et al. Opposing immune and genetic forces shape oncogenic programs in synovial sarcoma. bioRxiv. 2019 Jan 1;724302.

70. Ramilowski JA, Goldberg T, Harshbarger J, Kloppmann E, Lizio M, Satagopam VP, et al. A draft network of ligand–receptor-mediated multicellular signalling in human. Nature Communications. 2015 Jul 22;6:7866.

